# MYC Functions as a Switch for Natural Killer Cell-Mediated Immune Surveillance of Lymphoid Malignancies

**DOI:** 10.1101/503086

**Authors:** Srividya Swaminathan, Line D. Heftdal, Daniel F. Liefwalker, Renumathy Dhanasekaran, Anja Deutzmann, Crista Horton, Adriane Mosley, Mariola Liebersbach, Holden T. Maecker, Dean W. Felsher

**Affiliations:** Division of Oncology, Departments of Medicine and Pathology, Stanford University, Stanford, CA, USA; Division of Gastroenterology and Hepatology, Stanford University, Stanford, CA, USA; The Human Immune Monitoring Center (HIMC), Institute for Immunity, Transplantation and Infection, Stanford University School of Medicine, Stanford, California, USA

**Author notes:** **For correspondence**: Dean W. Felsher, MD PhD, Division of Medical Oncology, Departments of Medicine and Pathology, Stanford University, Stanford, CA.

## Abstract

The MYC oncogene drives T and B lymphoid malignancies, including Burkitt’s lymphoma (BL) and Acute Lymphoblastic Leukemia (ALL). Using CyTOF, we demonstrate a systemic reduction in natural killer (NK) cell-mediated surveillance in *SRα-tTA/Tet-O-MYC*^*ON*^ mice bearing MYC-driven T-lymphomas, due to an arrest in NK cell maturation. Inactivation of lymphoma-intrinsic MYC releases the brakes on NK maturation restoring NK homeostasis. Lymphoma-intrinsic MYC arrests NK maturation by transcriptionally repressing STAT1/2 and secretion of Type I Interferons (IFNs). Treating T-lymphoma-bearing mice with Type I IFN improves survival by rescuing NK cell maturation. In MYC-driven BL patients, low expression of both STAT1 and STAT2 correlates significantly with the absence of activated NK cells and predicts unfavorable clinical outcomes. Adoptive transfer of mature NK cells is sufficient to delay both T-lymphoma growth and recurrence post MYC inactivation. Our studies thus provide a rationale for developing NK cell-based therapies to effectively treat MYC-driven lymphomas in the future.

**Abbreviations:**

CyTOF: Cytometry Time of Flight
BL: Burkitt’s Lymphoma
ALL: Acute Lymphoblastic Leukemia
DLBCL: Diffuse Large B Cell Lymphoma
NK Cell: Natural Killer Cell
STAT1/2: Signal Transducer and Activator of Transcription 1/2
IFN: Interferon
DC: Dendritic Cell
MYC: v-myc avian myelocytomatosis viral oncogene homolog
PRECOG: Prediction of Clinical Outcomes from Genomic Profiles
BLI: Bioluminescence Imaging
TSS: Transcriptional Start Site
HTLV: Human T-lymphotropic Virus
hMYC: human MYC

---

Oncogene addiction is the phenomenon where a cancer becomes causally dependent on a single ‘driver’ oncogene for the maintenance of its proliferation and survival ^1,2^. Inhibition of the driver oncogene forms the basis of oncogene targeted therapy ^1–3^. Hence, a potential Achilles’ heel of human malignancies lies in their addiction to oncogenes such as *MYC, RAS* and *BCR-ABL1* ^4,5^. Targeted therapy has been successful against BCR-ABL1 (Philadelphia/Ph^+^)-induced leukemia ^6,7^, while therapies that successfully target MYC or RAS remain to be developed.

In order to develop targeted therapies for MYC-driven cancers, it is vital to understand whether MYC regulates both cell autonomous and non-autonomous processes, including host immunity. Oncogene addiction was previously assumed to be cell autonomous and immune-independent ^8^. Many recent studies illustrate that MYC and other oncogenes alone or cooperatively may regulate the tumor microenvironment and host immune responses in multiple tumor types ^9–16^. In particular, two reports highlighted that oncogenes may cooperate to more globally regulate the host immune system ^12,16^.

We have used a particularly tractable approach for studying the role of the host immune system during MYC-driven tumorigenesis through tetracycline (tet)-system regulated transgenic mouse models of cancer. Our inducible mouse models enable us to study how MYC inactivation elicits tumor regression through both cancer-intrinsic and cancer-extrinsic host immune-dependent mechanisms ^11,13^. CD4^+^ T cells appear to be essential for the sustained regression of cancers upon MYC inactivation ^11,13^. However, the changes in immune landscape during primary MYC-induced tumorigenesis in these models remain to be delineated.

Our goal was to delineate the global immunological changes resulting from primary overt MYC-driven lymphomagenesis, to identify anti-tumor immune subsets that can be developed as therapies to treat MYC-driven lymphomas. We hypothesized that oncogenic MYC directly perturbs the global immune composition at sites of primary lymphomagenesis to evade immune surveillance. Hence, using mass cytometry (CyTOF), we examined specific changes in the host immune composition both upon MYC activation as well as after subsequent MYC inactivation *in situ* in *SRα-tTA/tet-O-MYC* mice predisposed to developing autochthonous MYC-driven T cell lymphoblastic lymphoma ^1^.

We observed that MYC markedly suppresses maturation and production of natural killer (NK) cells in the lymphoma microenvironment. Furthermore, we found that NK cells play a key anti-tumorigenic role by delaying the growth and recurrence of MYC-driven T cell lymphomas, and are hence, suppressed by MYC during overt lymphomagenesis. We found a direct signaling mechanism by which lymphoma-intrinsic MYC suppresses NK cell-mediated immune surveillance. Our results provide a rationale for developing NK cell-based therapies that can be combined with MYC inhibitors to treat MYC-driven lymphomas.

## Results

### MYC-driven lymphomas exhibit disrupted splenic architectures

First, we evaluated the gross changes in lymphoid organs resulting from primary overt MYC-driven T cell lymphomagenesis in *SRα-tTA/tet-O-MYC* mice ^1^. Of note, SR*α* (a component of human T-lymphotropic virus; HTLV) restricts the overexpression of the human MYC (hMYC) transgene to the T cell lineage in *SRα-tTA/tet-O-MYC* mice, thus giving rise to systemically disseminated T cell lymphomas ^1^. Lymphoma-bearing mice displayed splenic germinal center disruption while inhibition of MYC partially rescued the germinal center architecture of the spleen (**Supplementary Figure 1**). These findings suggested that MYC overexpression in T cell lymphoma might remodel the splenic immune architecture.

### Oncogenic MYC perturbs the relative proportions of splenic immune subsets

Inducible regulation of the hMYC transgene specifically in malignant T-lymphocytes enables us to elucidate how lymphoma-intrinsic MYC impacts normal immune cells during primary lymphomagenesis. Our goal was to identify anti-tumor immune subsets that can be developed as therapies against MYC-driven lymphomas. Hence, using CyTOF ^17^, we delineated the global changes in the immune compositions of lymphoid organs during primary MYC-driven lymphomagenesis in *SRα-tTA/tet-O-MYC* mice.

Spleen was chosen for CyTOF experiments for two reasons. First, the spleen contains a diverse immune population including T and B lymphocytes, NK cells, dendritic cells (DCs), and myeloid cells that enable us to study interactions between lymphoma cells and their immune microenvironment *in situ*. Second, the lack of immature pre-T cells in the spleen allows us to visualize the infiltration of immature T cell lymphoblasts in MYC^ON^ mice and distinguish them from healthy and MYC^OFF^ mice. Primary splenic samples derived from overt lymphoma-bearing *SRα-tTA/tet-O-MYC* mice before (MYC^ON^, n = 9) and after (MYC^OFF^, 96h, n = 10) MYC inactivation were examined for immune subsets; and compared to age-matched normal wildtype spleens (Normal, n = 11).

Normal spleens contain mature single positive (SP) helper CD4^+^ T (CD3^+^ CD4^+^ CD8^-^) and cytotoxic CD8^+^ T (CD3^+^ CD4^-^ CD8^+^) cells, and are nearly devoid of immature double positive (DP, CD3^+^ CD4^+^ CD8^+^) and immature double negative (DN, CD3^+^ CD4^-^ CD8^-^) subsets. Consistent with the phenotype of a T cell lymphoma, the distribution of T cell subsets was altered in the MYC^ON^ cohort with an increase in the percentages of immature CD4^+^ CD8^+^ DP CD3^+^ T-cell leukemic blasts as compared to normal mice (Figures 1a-c, **Supplementary Figures 2a-b**). MYC inactivation (MYC^OFF^) resulted in elimination of most DP T cell blasts and restored the distribution of splenic T cell subsets to normal levels, demonstrating MYC-addiction, as previously described ^1^ (Figures 1a-c, **Supplementary Figures 2a-b**). Malignant CD3^+^ T-lymphocytes in MYC^ON^ mice are TCR*α*β^+^, thus leading to a reduction in the percentages of TCRγδ^+^ T cells (**Supplementary Figures 2c-d**).

We next examined the other non-malignant immune compartments that do not express the T cell specific hMYC transgene in *SRα-tTA/tet-O-MYC* mice before (MYC^ON^) and after (MYC^OFF^) MYC inactivation, and compared these to normal mice. Amongst these normal subsets, percentages of NK (CD3^-^ NKp46^+^, Figure 1d, **Supplementary Figures 2e-f**), and B cell (CD19^+^, Figure 1e, **Supplementary Figures 3a-b**) subsets were significantly lowered in MYC^ON^ overt lymphoma-bearing mice. Yet, NK (Figure 1d, **Supplementary Figures 2e-f**), and B cell (Figure 1e, **Supplementary Figures 3a-b**) percentages were restored close to normal levels in MYC^OFF^ mice. The relative proportions of other immune compartments including myeloid compartments such as dendritic cells (DCs) and neutrophils were unaltered by the activation of oncogenic MYC (**Supplementary Figures 3c-f**).

**Figure 1:**
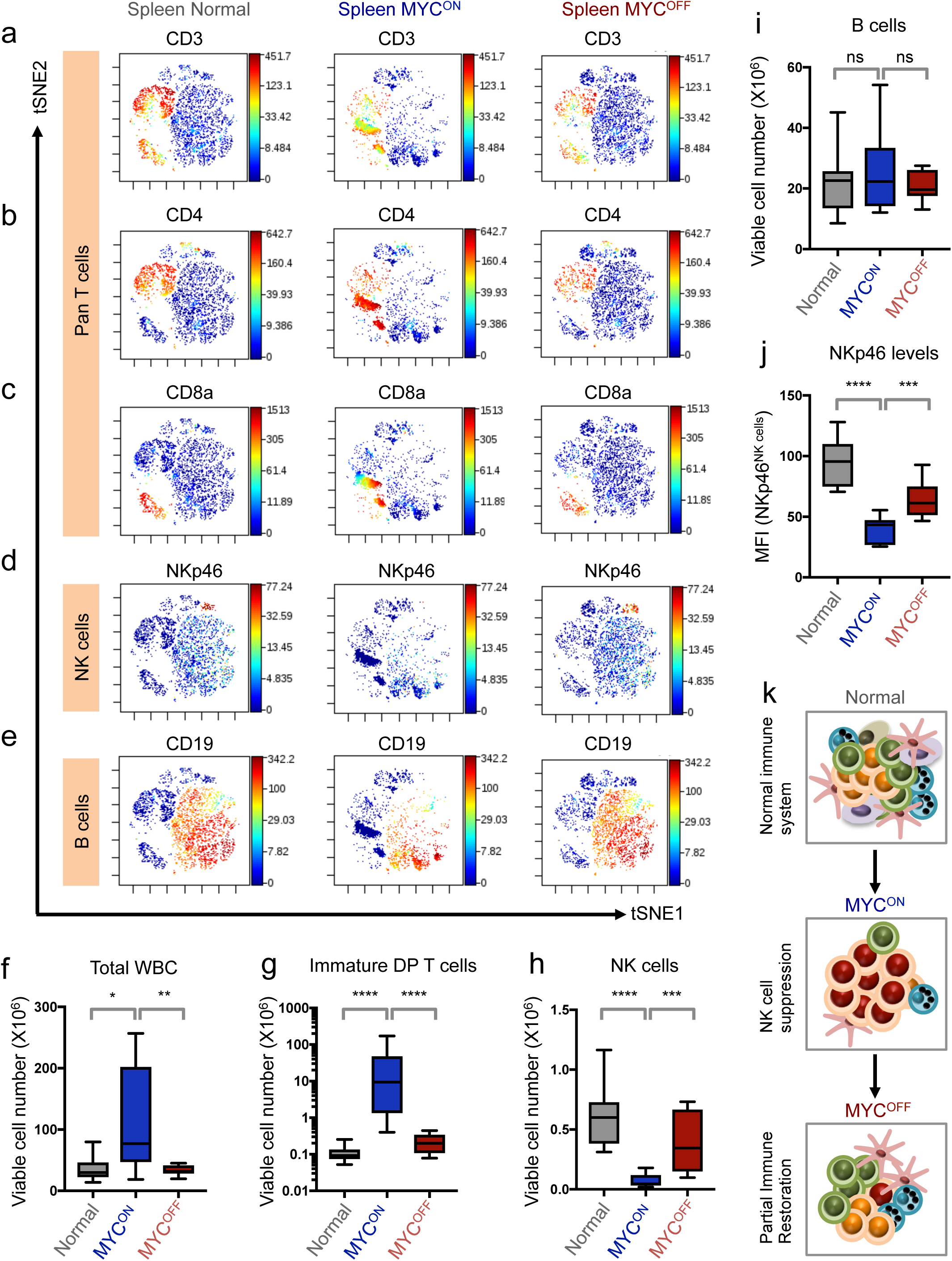
NK cells are reduced in numbers and maturation in primary MYC-driven T cell lymphomas. **(a-e)** viSNE of CyTOF data depicting splenic CD3^+^ T (**a**), splenic CD4^+^ T **(b)**, and CD8a^+^ T **(c)**, NKp46^+^ NK (**d**), and CD19^+^ B (**e**) cell compositions of one representative mouse from normal (n = 11), SR*α*-tTA MYC^ON^ (n = 9), and SR*α*-tTA MYC^OFF^ (doxycycline 96 h, n = 10) groups. **(f-i)** Quantification of absolute counts of total WBC (**f**), immature CD3^+^ CD8^+^ CD4^+^ DP lymphoma T cells (**g**), CD3^-^ NKp46^+^ mature NK (**h**), and CD19^+^ B (**i**) cells in normal (n = 11), SR*α*-tTA MYC^ON^ (n = 9), and SR*α*-tTA MYC^OFF^ (doxycycline 96 h, n = 10) mice subjected to CyTOF. (**j**) Median fluorescence intensity (MFI) of NKp46 on splenic NK cells from normal (n = 11), SR*α*-tTA MYC^ON^ (n = 9), and SR*α*-tTA MYC^OFF^ (doxycycline 96 h, n = 10) mice. **(k)** Schematic depicting the reversible phenomenon of NK cell (blue) suppression in MYC-driven lymphomas (red). P-values have been calculated using the Mann-Whitney test. P-values: ns = not significant; *p < 0.05; **p < 0.01; ***p < 0.001; ****p < 0.0001.

### Immune composition of MYC-driven lymphomas is altered independently of splenomegaly

Lymphomagenesis is often associated with splenomegaly and increased splenic cellularity. Therefore, we had to rule out the possibility that the apparent reduction in immune subsets such as, NK and B cells was because they were being passively outnumbered by invading MYC-driven malignant lymphocytes. To address this, we measured the absolute cell numbers of the immune subsets in the splenic samples evaluated by CyTOF. Notably, we observed a statistically significant increase in the cellularity of lymphoma spleens (Figure 1f). As expected, the absolute cell counts of CD3^+^ pan T (**Supplementary Figure 4a**), TCR*α*β^+^ CD3^+^ T (**Supplementary Figure 4b**) and immature CD4^+^ CD8^+^ double positive (DP) CD3^+^ leukemic T cells (Figure 1g) were increased in MYC^ON^ mice when compared to normal and MYC^OFF^ mice. Characteristic with the properties of a lymphoma, abundant multiplication of the TCR*α*β^+^ CD3^+^ T cell lymphoblasts was accompanied by a significant reduction in numbers of CD3^+^ TCRγδ^+^ γδ T cells in MYC^ON^ mice (**Supplementary Figure 4c**).

Oncogenic MYC significantly lowered absolute cell numbers of CD3^-^ NKp46^+^ NK cells, whereas MYC inhibition significantly reversed this effect (Figure 1h). Absolute numbers of B cells were unaltered in MYC-driven lymphomas as compared to healthy controls and MYC^OFF^ mice (Figure 1i), although B cell percentages had been significantly reduced in lymphoma-bearing mice (Figure 1e, **Supplementary Figures 3a-b**). MYC-driven lymphomagenesis significantly increased numbers of pan CD11c^+^ DCs and neutrophils (**Supplementary Figures 4d-e**). The increase in numbers of DCs and neutrophils during MYC-driven lymphomagenesis corroborated the previously well-characterized pro-tumorigenic functions of these subsets in lymphomas ^18–20^.

### NK cells represent a key anti-tumorigenic immune compartment suppressed during MYC-driven lymphomagenesis

Our goal was to identify anti-tumor immune subsets that can be developed as therapies against MYC-driven lymphomas. Amongst the anti-tumorigenic immune subsets (NK and B cells) altered during MYC-driven lymphomagenesis, NK cells particularly stood out for the following reasons: First, NK cells were the sole subset significantly altered both in absolute cell numbers (Figure 1h) and percentages (Figure 1d, **Supplementary Figures 2e-f**). Second, MYC inactivation restores NK cell compositions suggesting that oncogenic MYC may directly regulate this immune subset (Figures 1d, h). Third, we observed a significant suppression of maturation and cytotoxicity marker NKp46 ^21^ on the surface of residual NK cells in MYC^ON^ spleens when compared to normal and MYC^OFF^ spleens (Figure 1j). Fourth, NK cells are attractive as a potential cell-based immunotherapy against MYC-driven lymphomas. Hence, we focused specifically on how oncogenic MYC alters the NK subset during lymphomagenesis.

We confirmed our CyTOF results which show an inverse correlation between the activity of MYC and NK cell presence, by measuring NK cell compositions before and after MYC inactivation by conventional flow cytometry (**Supplementary Figure 5**). Next, using CIBERSORT ^22^, a computational method for determining immune compositions from bulk gene expression data, we estimated changes in NK composition in CyTOF-matched spleens from normal (n = 3), SR*α-*tTA MYC^ON^ (lymphoma, n = 3) and SR*α-*tTA MYC^OFF^ (regressed lymphoma, n = 3) mice. Concordant with CyTOF results, CIBERSORT also demonstrates a reduction in the relative proportion of the NK cells in MYC^ON^ mice when compared to normal and MYC^OFF^ mice (**Supplementary Figure 6**). Hence, using three different platforms (CyTOF, flow cytometry, and CIBERSORT), we were able to robustly demonstrate that oncogenic MYC causally and reversibly reduces splenic NK cell-mediated immune surveillance during lymphomagenesis (Figure 1k).

### NK cell suppression during MYC-driven lymphomagenesis is systemic in nature

We examined whether the phenomenon of NK cell suppression occurs in other lymphoid organs infiltrated by malignant T cell lymphoblasts, including blood (Figures 2a-e) and bone marrow (**Supplementary Figure 7**). Circulating NK cell percentages (Figures 2b-c), numbers per µl of blood (Figure 2d), and maturation (NKp46 MFI, Figure 2e) were significantly reduced in overt lymphoma mice (MYC^ON^, n = 6), as compared to normal (n = 6) and MYC-inactivated (MYC^OFF^, n = 6) mice.

**Figure 2:**
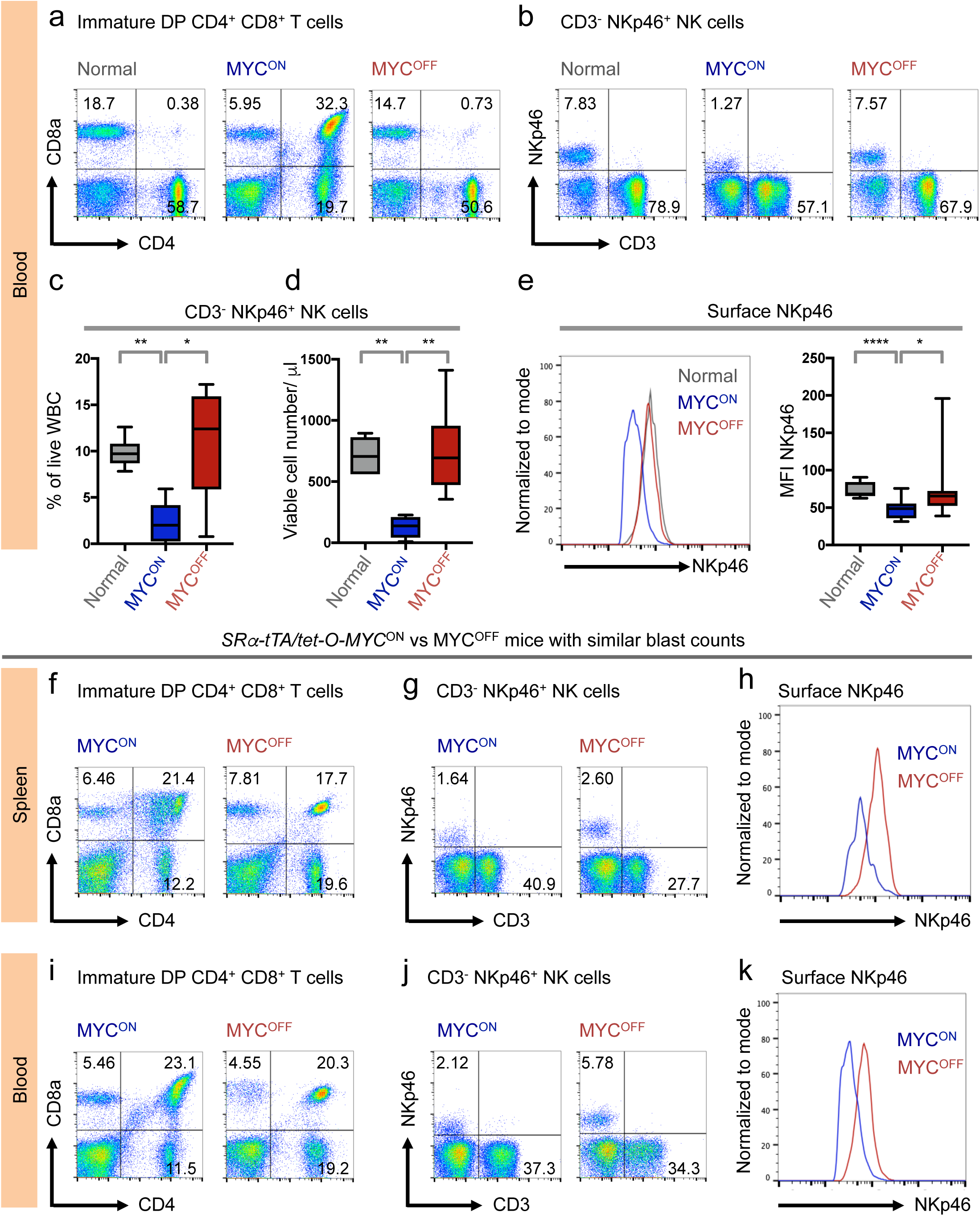
NK cell suppression in MYC-driven lymphomas is systemic and depends on MYC status. **(a-b)** Representative flow cytometry plots depicting percentages of circulating immature CD4^+^ CD8^+^ T (**a**), and CD3^-^ NKp46^+^ NK (**b**) cells in normal (n = 6), SR*α*-tTA MYC^ON^ (n = 6), and SR*α*-tTA MYC^OFF^ (doxycycline 96 h, n = 6) mice. **(c-d)** Quantification of percentages (**c**), and absolute counts per µl of blood (**d**) of CD3^-^ NKp46^+^ NK cells, in normal (n = 6), SR*α*-tTA MYC^ON^ (n = 6), and SR*α*-tTA MYC^OFF^ (doxycycline 96 h, n = 6) mice. **(e)** Comparison of MFI of surface NKp46 on circulating NK cells in normal (n = 6), SR*α*-tTA MYC^ON^ (n = 6), and SR*α*-tTA MYC^OFF^ (doxycycline 96 h, n = 6) mice. **(f-g)** Representative flow cytometry plots depicting percentages of splenic immature CD4^+^ CD8^+^ T (**f**), and CD3^-^ NKp46^+^ NK (**g**) cells in SR*α*-tTA MYC^ON^ and SR*α*-tTA MYC^OFF^ (doxycycline 96 h) mice with comparable blast counts. **(h)** Comparison of surface NKp46 levels (MFI) on splenic NK cells from SR*α*-tTA MYC^ON^ and SR*α*-tTA MYC^OFF^ (doxycycline 96 h) mice with comparable blast counts. **(i-j)** Representative flow cytometry plots depicting percentages of circulating immature CD4^+^ CD8^+^ T (**i**), and CD3^-^ NKp46^+^ NK (**j**) cells in SR*α*-tTA MYC^ON^ and SR*α*-tTA MYC^OFF^ (doxycycline 96 h) mice with comparable blast counts. **(k)** Comparison of surface NKp46 levels (MFI) on circulating NK cells from SR*α*-tTA MYC^ON^ and SR*α*-tTA MYC^OFF^ (doxycycline 96 h) mice with comparable blast counts. P-values have been calculated using the Mann-Whitney test. P-values: ns = not significant; *p < 0.05; **p < 0.01; ****p < 0.0001.

Next, we investigated whether MYC-driven T cell lymphomagenesis leads to NK cell suppression in the bone marrow. T cell lymphoma in our model manifests with significant bone marrow involvement (**Supplementary Figures 7a-d**). CD3^-^ NKp46^+^ mature NK cells were significantly suppressed in bone marrow of lymphoma-bearing mice (MYC^ON^, n = 7), as compared to normal (n = 9) and MYC-inactivated (MYC^OFF^, n = 7) mice (**Supplementary Figures 7e-f**). NK cell numbers in the bone marrow were significantly restored following MYC inactivation (**Supplementary Figures 7e**).

### NK cell suppression during lymphomagenesis is MYC-dependent

We next demonstrated the dependency of NK suppression on MYC status, by comparing percentages and surface NKp46 expression of splenic (Figures 2f-h) and circulating (Figures 2i-k) NK cells in MYC^OFF^ mice (with residual blasts) alongside lymphoma-bearing mice (MYC^ON^) with similar blast percentages. The presence of residual lymphoblasts in MYC^OFF^ mice may be attributed to (**1**) initial high disease burden before inactivating MYC, or (**2**) partial differentiation of lymphoblasts after MYC inactivation ^23^. We observed that MYC inactivation increases NK cell percentages (Figures 2g, j) and the expression of NK cell maturation and cytotoxicity marker NKp46 (MFI surface NKp46, Figures 2h, k) even in the presence of residual lymphoblasts (Figures 2f, i). Hence, we infer that NK cell suppression is MYC-dependent, and does not occur simply due to the creation of niche space vacated by the clearance of lymphoma cells following MYC inactivation.

### Maturation of NK cells is arrested during MYC-driven lymphomagenesis

We investigated whether reduction in mature CD3^-^ NKp46^+^ NK cells during MYC-induced lymphomagenesis occurs because of increased cell death. Surprisingly, we observed reduced death of NK cells at sites of lymphomagenesis in lymphoma-bearing MYC^ON^ mice, in comparison to normal and MYC^OFF^ cohorts (**Supplementary Figure 8**). Hence, suppression of NK subset during lymphomagenesis is not caused by increased apoptosis of NK cells induced by lymphomagenic MYC.

The disappearance of NK cells from the bone marrow of MYC^ON^ mice (**Supplementary Figure 7**) suggested that MYC-driven lymphomagenesis might abrogate the replenishment of mature CD3^-^NKp46^+^ NK cells in the periphery possibly by blocking early NK cell development ^24–27^. We tested this possibility by comparing the absolute counts of NK cell precursors (NKP, CD122^+^ NKp46^-^, Figure 3a), and the more differentiated CD122^+^ NKp46^+^ iNK (immature NK, Figure 3b) cells in bone marrow from normal (n = 8), SR*α-*tTA MYC^ON^ (lymphoma, n = 8), and SR*α-*tTA MYC^OFF^ (regressed lymphoma, n = 8) mice. While we observed no significant changes in the numbers of NKP across the three groups, the numbers of iNK cells were significantly reduced in MYC^ON^ bone marrow, in comparison to MYC^OFF^ and normal bone marrows (Figures 3a-b). Consistent with these observations, the ratio of counts of NKP vs iNK were significantly enhanced in individual MYC^ON^ mice in comparison to healthy and MYC^OFF^ mice (Figure 3c).

**Figure 3:**
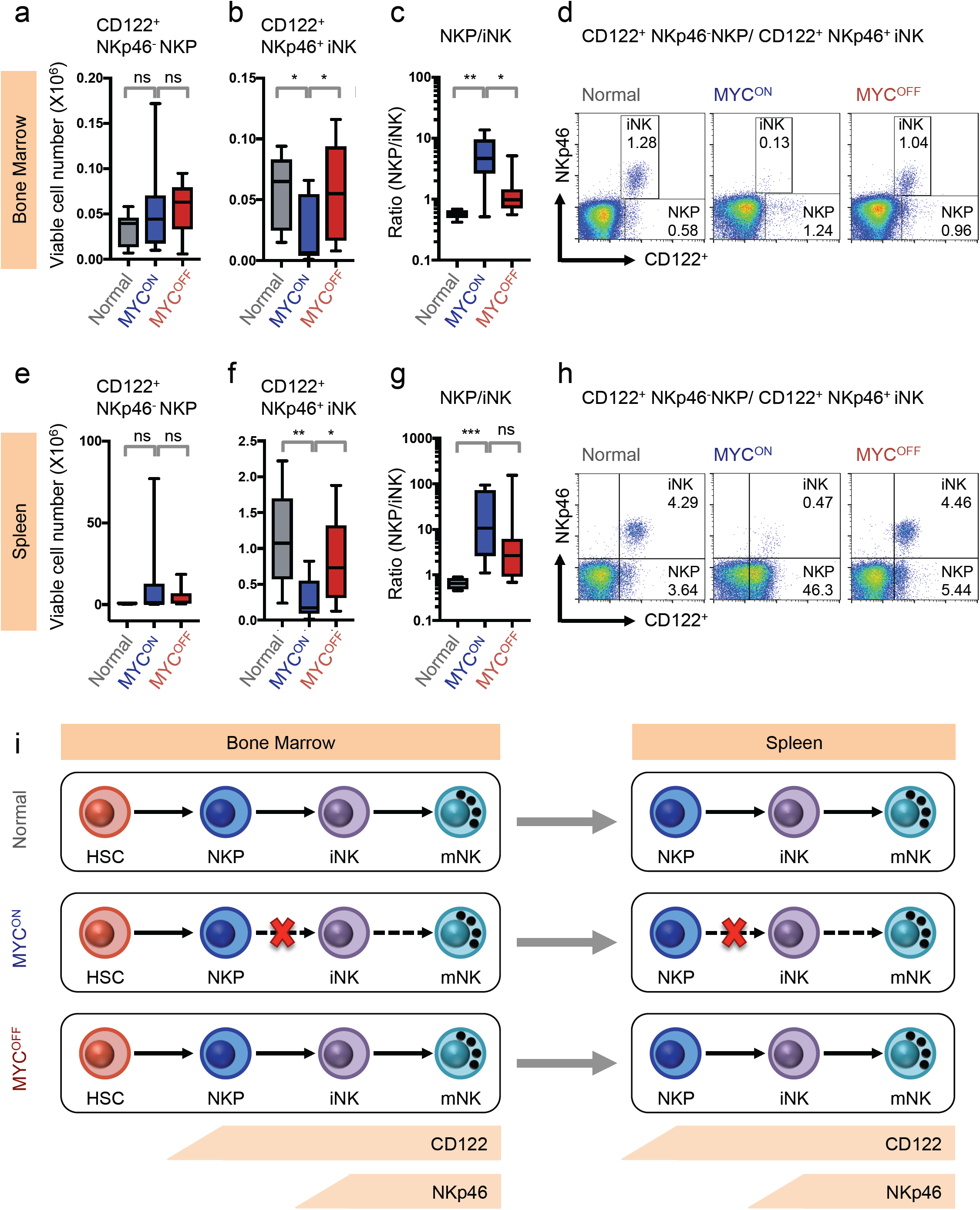
Maturation of NK cells is arrested in mice bearing overt MYC-driven T cell lymphomas. **(a-b)** Quantification of viable cell numbers of NK precursor cells (NKP, CD122^+^ NKp46^-^, (**a**)), and immature NK cells (iNK, CD122^+^ NKp46^+^, (**b**)) in normal (n = 8), SR*α*-tTA MYC^ON^ (n = 8), and SR*α*-tTA MYC^OFF^ (doxycycline 96 h, n = 8) bone marrow measured by flow cytometry. **(c)** Comparison of proportions of numbers of NKP and iNK cells (NKP/iNK) between normal (n = 8), SR*α*-tTA MYC^ON^ (n = 8), and SR*α*-tTA MYC^OFF^ (doxycycline 96 h, n = 8) bone marrow. **(d)** Representative flow cytometry plots depicting NK precursor cells (NKP, CD122^+^ NKp46^-^), and immature NK cells (iNK, CD122^+^ NKp46^+^) in normal (n = 8), SR*α*-tTA MYC^ON^ (n = 8), and SR*α*-tTA MYC^OFF^ (doxycycline 96 h, n = 8) bone marrow. **(e-f)** Quantification of viable cell numbers of NK precursor cells (NKP, CD122^+^ NKp46^-^, (**d**)), and immature NK cells (iNK, CD122^+^ NKp46^+^, (**e**)) in normal (n = 8), SR*α*-tTA MYC^ON^ (n = 8), and SR*α*-tTA MYC^OFF^ (doxycycline 96 h, n = 8) spleens measured by flow cytometry. **(g)** Comparison of proportions of numbers of NKP and iNK cells (NKP/iNK) between normal (n = 8), SR*α*-tTA MYC^ON^ (n = 8), and SR*α*-tTA MYC^OFF^ (doxycycline 96 h, n = 8) spleens. **(h)** Representative flow cytometry plots depicting NK precursor cells (NKP, CD122^+^ NKp46^-^), and immature NK cells (iNK, CD122^+^ NKp46^+^) in normal (n = 8), SR*α*-tTA MYC^ON^ (n = 8), and SR*α*-tTA MYC^OFF^ (doxycycline 96 h, n = 8) bone marrow. **(i)** Model depicting the arrest in differentiation from NK precursor (NKP) stage to immature NK (iNK) stage during MYC-driven lymphomagenesis. Arrested step in NK cell differentiation is indicated by a red cross. Dashed arrows indicate stages of NK cell differentiation that do not occur because of the arrest. All p values have been calculated using Mann-Whitney test. P-values: ns = not significant; *p < 0.05; **p < 0.01; ***p < 0.001.

We infer that oncogenic MYC blocks the expression of NKp46 and hence, the transition from NKP to iNK stage of NK cell maturation in the bone marrow ^24,25^. The spleens of MYC^ON^ mice exhibited no changes in the CD122^+^ NKp46^-^ NKP, and a significant reduction in the more differentiated CD122^+^ NKp46^+^ iNK fraction (Figures 3e-h), concordant with the changes observed in their corresponding bone marrows (Figures 3a-d). Hence, NK cells are systemically suppressed during MYC-induced lymphomagenesis because of arrested NK cell maturation in the bone marrow that translates to the periphery (Figure 3i).

### Suppression of Type I IFN signature during primary MYC-driven lymphomagenesis may block NK cell-mediated immune surveillance

We next investigated the molecular mechanisms by which oncogenic MYC may suppress NK cell-mediated immune surveillance during primary T and B cell lymphomagenesis. To this end, we analyzed the global transcriptomic profiles of bulk splenic cells from normal (n = 3), SR*α-*tTA MYC^ON^ (lymphoma, n = 3) and SR*α-*tTA MYC^OFF^ (regressed lymphoma, n = 3) mice (GSE106078). We observed that the expression profiles of MYC^OFF^ spleens clustered closely with normal spleens, thus confirming that transcriptional changes induced by MYC were largely reversible upon its inactivation (Figure 4a). Gene set enrichment analyses (GSEA) revealed that the JAKs (Janus Kinases) – STAT1/2 (Signal Transducer and Activator of Transcription 1/2) – Type I IFN (Interferon) pathway was significantly suppressed in MYC^ON^ mice when compared to normal and MYC^OFF^ mice (Figures 4b-d, **Supplementary Figure 9**). Of note, the activation of STAT1/2 post MYC inactivation resembles an anti-viral immune response that is associated with production of Type I IFNs and subsequent activation of NK cell-mediated immune surveillance ^28^.

STAT1/2-Type I IFN signaling has been previously demonstrated to be required for maturation and expansion of NK cells before the expression of NKp46 at the iNK stage ^29–35^. Of note, mice deficient in Type I IFN receptor (*IFNαR*^-/-^) show significantly lower numbers of NK cells in spleen when compared to normal mice ^29,35^; a phenomenon we observe during MYC-driven lymphomagenesis (Figure 1h). Concordant with our results of NK cell suppression as a result of MYC-driven lymphomagenesis (Figures 1-2), we observed transcriptional repression of a number of Type I IFN regulated genes that are specific to NK-mediated immune surveillance in MYC^ON^ mice, when compared to normal and MYC^OFF^ mice (Figure 4e). These findings suggested the possibility of existence of a link between Type I IFN signature and the observed splenic NK cell profiles of each group.

**Figure 4:**
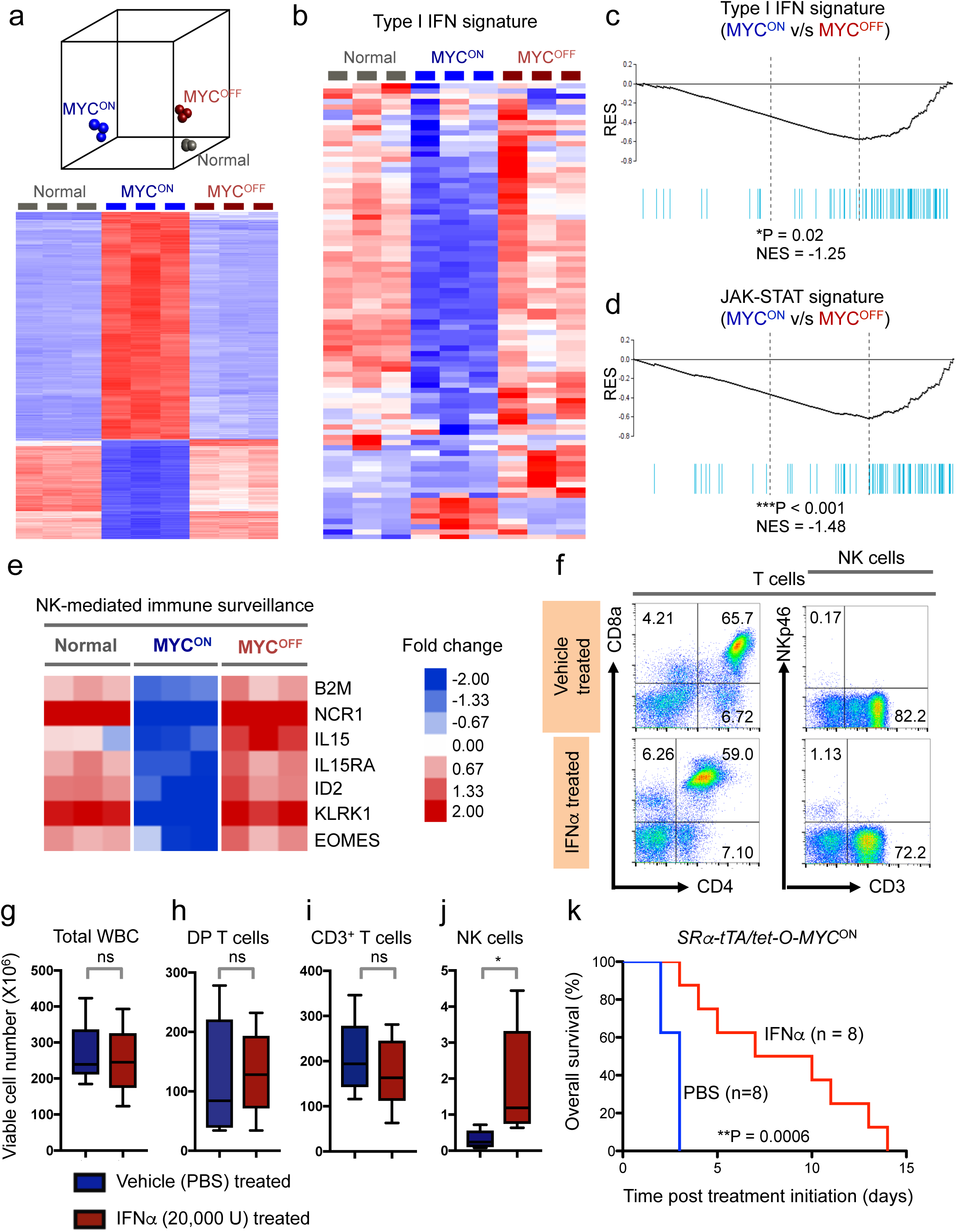
Escape from NK surveillance during MYC-driven lymphomagenesis is mediated by suppression of Type I IFN signaling. **(a)** Principal component analysis (PCA, top) and heatmap (bottom) depicting global transcriptomic profiles of bulk splenic samples derived from normal (n = 3), SR*α*-tTA MYC^ON^ (n = 3) and SR*α*-tTA MYC^OFF^ (n = 3) mice, obtained by RNA sequencing (GSE106078). **(b)** Heatmap showing suppression of Type I IFN signaling in spleens of SR*α*-tTA MYC^ON^ (n = 3) mice when compared to normal (n =3) and SR*α*-tTA MYC^OFF^ (n = 3) mice. **(c)** Gene Set Enrichment Analysis (GSEA) depicting suppression of Type I IFN signature in spleens of SR*α*-tTA MYC^ON^ (n = 3) mice when compared to SR*α*-tTA MYC^OFF^ (n =3) mice. **(d)** GSEA depicting suppression of JAK-STAT signature in spleens of SR*α*-tTA MYC^ON^ (n = 3) mice when compared to SR*α*-tTA MYC^OFF^ (n =3) mice. **(e)** Heatmap comparing expression of key NK cell genes in splenic samples from normal (n = 3), SR*α*-tTA MYC^ON^ (n = 3) and SR*α*-tTA MYC^OFF^ (n = 3) mice. **(f)** Representative flow cytometry plots comparing percentages of CD4^+^ CD8^+^ DP T (left) and CD3^-^ NKp46^+^ NK (right) cells between *SRα-tTA/tet-O-MYC* mice in IFNα-treated and vehicle (PBS)-treated groups for three days. **(g-j)** Quantification of absolute counts of total WBC (**g**), CD4^+^ CD8^+^ DP T (**h**), CD3^+^ pan T (**i**), and CD3^-^ NKp46^+^ NK cells (**j**) in *SRα-tTA/tet-O-MYC* mice bearing overt lymphomas (SR*α*-tTA MYC^ON^) treated for three days with either IFNα (n = 5), or vehicle (PBS, n = 5). **(k)** Comparison of overall survival (OS) probabilities between PBS (n = 7) and IFN*α* (n = 7) -treated *SRα-tTA/tet-O-MYC* mice bearing overt lymphomas (SR*α*-tTA MYC^ON^, age = 2-3 months). P-values have been calculated using Mann-Whitney test. ns = not significant, *p < 0.05, **p < 0.01.

Next, we compared transcript levels of STAT1, STAT2 and other important Type I IFN signaling components in normal B cells (MYC^LOW^, n = 4), pre-lymphoma B cells from Eµ-MYC mice (MYC^HIGH^, n = 4), and overt B cell lymphoma from Eµ-MYC mice (MYC^HIGH^, n = 2), which is an independent mouse model of MYC-driven B cell lymphoma (GSE51011, ^36^). Sequential increase in MYC during stepwise B cell transformation is accompanied by transcriptional suppression of STAT1/2-Type I IFN signature (**Supplementary Figure 10**). Hence, suppression of STAT1/2-Type I IFN signaling is a signature of both T cell and B cell MYC-driven lymphomagenesis.

### MYC-mediated suppression of NK cells can reversed by Type I IFN treatment

We investigated whether short-term (3 days) administration of Type I IFN to SR*α-*tTA MYC^ON^ mice bearing overt T cell lymphoma can rescue MYC-mediated NK cell suppression, and increase infiltration of NK cells into spleens (Figures 4f-j). The percentages and numbers of CD3^-^ NKp46^+^ NK cells were significantly increased in IFN*α*-treated mice (n = 5) when compared to corresponding litter-matched lymphoma mice treated with vehicle (PBS, n = 5) (Figures 4f, 4j). Of note, total numbers and percentages of WBC, CD3^+^ T cells and DP immature T cells remain unaltered between the control and IFN*α*-treated groups (Figures 4g-i), thus highlighting that Type I IFNs specifically regulate NK cell homeostasis and do not exert any direct anti-proliferative effects on tumor cells. Long-term administration of Type I IFNs to SR*α-* tTA MYC^ON^ mice bearing overt T cell lymphomas (n = 8) improves overall survival (OS) in comparison to vehicle (PBS)-treated (n = 8) lymphoma-bearing mice (Figure 4k). Hence, we conclude that MYC-driven lymphomagenesis may subvert normal immunological processes of NK cell homeostasis by suppressing STAT1/2 signaling cascade and Type I IFN production.

### Lymphoma-intrinsic MYC suppresses Type I IFN production by tumor cells

We hypothesized that signals blocking NK cell maturation arise from MYC-driven lymphoma cells for two reasons. First, we observed that suppression of NK cells is dependent on the MYC status of T cell lymphoblasts (Figures 2f-k). Second, we showed that MYC-overexpressing lymphoblasts display suppression of STAT1/2-Type I IFN signaling during primary lymphomagenesis (**Supplementary Figure 10**). Hence, as the next step, we investigated whether modulation of oncogenic MYC *in vitro* in MYC-driven B and T lymphoma cell lines differentially regulates STAT1/2-Type I IFN signaling.

Since BL is a classic human MYC-associated lymphoid cancer, we measured signaling changes by Phospho-CyTOF before (MYC^ON^) and after (MYC^OFF^) MYC inactivation in an EBV-transformed human B cell line that expresses MYC regulated by the Tet System (P493-6), and mimics BL ^37^ (**Supplementary Figure 11**). We confirmed that MYC is inducibly expressed in P493-6 cells (Figure 5a). Amongst the key immune regulatory cytokine signaling pathways that operate through the JAKs and STATs, MYC suppressed the activation of STAT1 in P493-6 cells (Figures 5b-c, **Supplementary Figure 11**). MYC inactivation increased STAT1 phosphorylation and activity (Figures 5b-c), STAT1 transcription (Figure 5d, **Supplementary Figure 12a**) and STAT1 protein production (Figure 5c). Transcription of STAT2, the binding partner of STAT1, was also induced post MYC inactivation in P493-6 cells (Figure 5d, **Supplementary Figure 12b**). Indeed, the transcript level of Type I IFN, IFN*α*2 was significantly elevated after MYC inactivation in P493-6 cells (MYC^OFF^, Figure 5e). Comprehensive cytokine profiling of the P493-6 cells pre-and post-MYC inactivation (**Supplementary Figure 13**) demonstrated that the secretion of IFN*α* is significantly upregulated in the MYC^OFF^ cells in comparison to the MYC^ON/HIGH^ population (Figure 5f).

Tumor-intrinsic signaling changes post MYC inactivation in our T cell lymphoma line derived from *SRα-tTA/tet-O-MYC* mice (**Supplementary Figure 14**) mirrored those observed in the human BL-like model with MYC inactivation leading to activation of STAT1/2-Type I IFN signaling (**Supplementary Figure 15**). Our findings suggested that lymphoma-intrinsic MYC represses STAT1/2-Type I IFN signaling; thereby blocking the maturation of NK cells in the tumor microenvironment, and NK cell-mediated immune surveillance.

**Figure 5:**
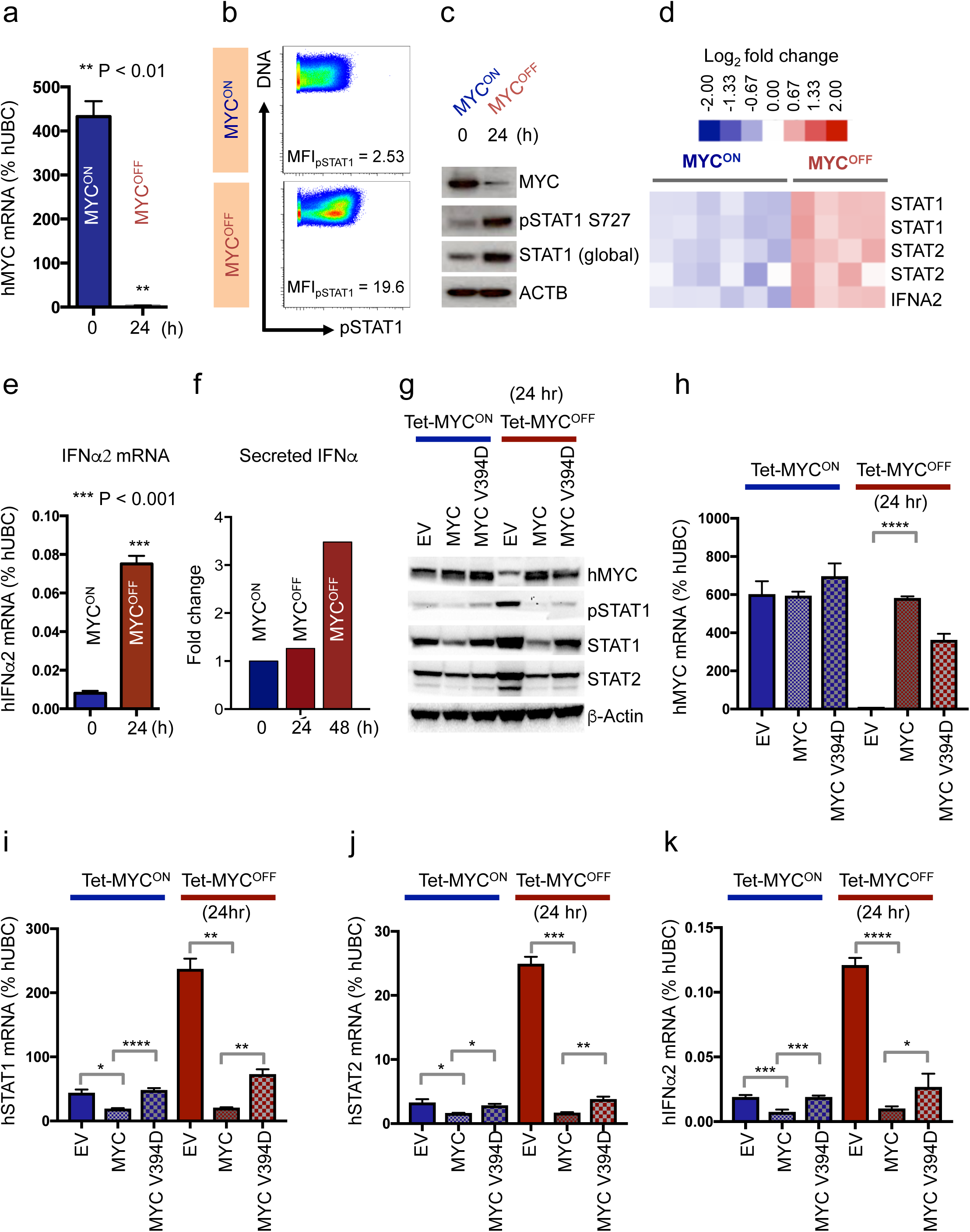
MYC transcriptionally represses the STAT1/2-Type I IFN signaling cascade associated with NK cell maturation. **(a)** Quantitative real time PCR for MYC in P493-6 BL human cells before and after MYC inhibition for 24h by doxycycline (n = 3, mean ± s.d). **(b)** Phospho mass cytometry (Phospho-CyTOF) depicting changes in pSTAT1 levels before (MYC^ON^) and after (MYC^OFF^) MYC inactivation for 24h in P493-6 cells. **(c)** Immunoblotting to validate changes in pSTAT1 observed by phospho-CyTOF. (**d**) Heat map depicting transcriptional changes in STAT1/2-Type I IFN signaling cascade before (MYC^ON^, n = 6) and after MYC inactivation (MYC^OFF^, 24h, n = 4) in P493-6 cells using expression arrays. **(e)** Quantitative real time PCR for Type I IFN*α*2 in P493-6 BL human cells before and after MYC inactivation for 24h by doxycycline (n = 3, mean ± s.d). **(f)** Luminex assay depicting changes in Type I IFN*α*2 secreted from P493-6 cells before and after MYC inactivation for 24 and 48h. **(g)** Immunoblotting in P493-6 BL cell lines expressing empty vector (EV), MYC overexpression vector (MYC), or a MYC mutant that fails to bind MIZ1 (MYC V394D), pre-(Tet-MYC^ON^) and post (Tet-MYC^OFF^) MYC inactivation. **(h-k)** Quantitative real time PCR for MYC (**h**), STAT1 (**i**), STAT2 (**j**), and Type I IFN*α*2 (**k**) in P493-6 BL cell lines expressing empty vector (EV), MYC overexpression vector (MYC), or a MYC mutant that fails to bind MIZ1 (MYC V394D), pre-(Tet-MYC^ON^) and post (Tet-MYC^OFF^) MYC inactivation (n = 3, mean ± s.d). All p values have been calculated using Student’s t-test. *p < 0.05; **p < 0.01; ***p < 0.001; ****p < 0.0001.

### MYC transcriptionally represses the activation of STAT1/2-Type I IFN signaling

We tested the possibility of MYC transcriptionally repressing the STAT1/2-Type I IFN signaling cascade. Although widely known as a transactivator, MYC can inhibit the transcription of genes by sequestering transactivators such as MIZ1 and Sp1/Sp3 ^38–40^, and binding to Initiator (Inr) elements in gene promoters. For example, MYC inactivation releases MYC-MAX heterodimers from the binding sites of tumor suppressor genes such as *Cdkn2b* and *Cdkn1b*, thereby promoting MIZ1-mediated transcriptional activation of these genes ^38–40^.

Concordant with a previous study where MYC was shown to transcriptionally repress STAT1 ^41^, we found that MYC binds the STAT1 promoter in P493-6 human BL and mouse MYC-driven T cell lymphoma (**Supplementary Figure 16**). In order to delineate the molecular mechanism behind MYC-mediated transcriptional repression of Type I IFN signaling, we compared levels of STAT1, STAT2, and IFN*α*2 in MYC-driven lymphoma cell lines overexpressing MYC, or a MYC mutant that fails to sequester MIZ1 (MYC V394D), or the corresponding empty vector (EV). While MYC overexpression in P493-6 cells effectively repressed the production of STAT1/2 (Figures 5g-j) and IFN*α*2 (Figure 5k), the MYCV394D mutant significantly rescued the activation of STAT1/2-Type I IFN signaling, both before and after inactivation of tetracycline-controlled MYC (Tet-MYC). Our results show that MYC transcriptionally represses STAT1/2-Type I IFN signaling partially by sequestering transactivators such as MIZ1. In MYC-driven T-ALL cells, we observe that MYC-mediated suppression of the Type I IFN signaling cascade is independent of MIZ1 (**Supplementary Figure 17**), suggesting the possible involvement of other MYC binding partners such as SP1/3 ^40^.

### MYC^High^ human lymphomas exhibit suppression of STAT1/2 and NK cell-mediated immune surveillance

We examined MYC-driven human lymphoma patients for evidence of repression of STAT1/2 signaling and NK cell-mediated immune surveillance. First, we analyzed human lymphomas that often exhibit MYC hyperactivation, such as Burkitt’s Lymphoma (BL) and Diffuse Large B Cell Lymphoma (DLBCL, GSE4475 ^42^). We observed that STAT1 and STAT2 levels inversely correlate with MYC levels in BL (n = 34, blue) and DLBCL (n = 187, orange) patients (Figures 6a-b). Of note, the BL subset driven directly by MYC translocations ^43^ displays the highest MYC levels in combination with the lowest levels of STAT1 and STAT2 (Figures 6a-b). Hence, we concluded that natural addiction to MYC in the BL subgroup might lead to the suppression of STAT1/2-Type I IFN signaling.

**Figure 6:**
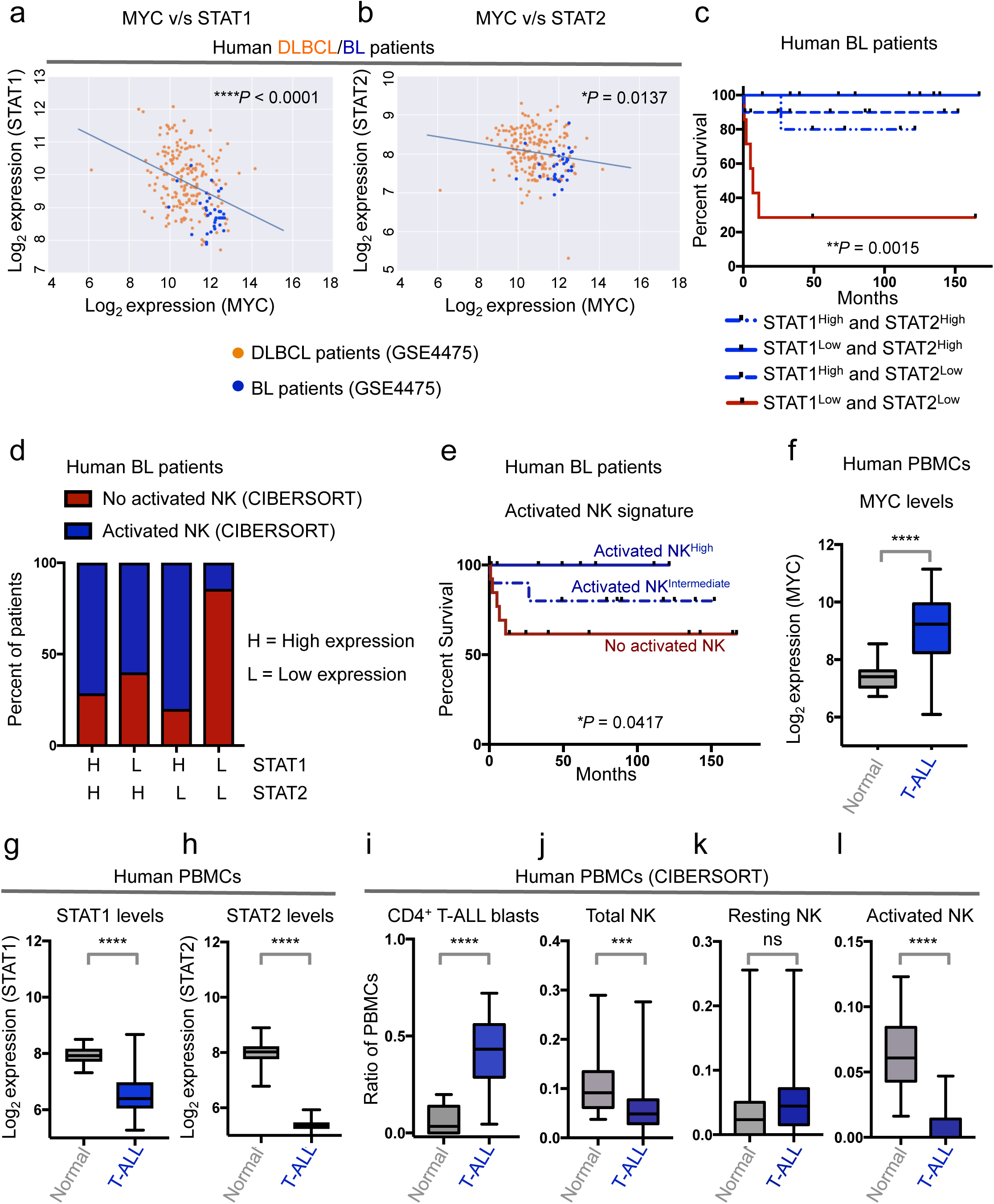
MYC^High^ human lymphomas exhibit decreased levels of STAT1/2 and NK cell-mediated immune surveillance. **(a-b)** Correlations between mRNA levels of MYC and STAT1 (**a**), or MYC and STAT2 (**b**) in classical MYC-driven human lymphoma patient samples (n = 187, DLBCL, orange; n = 34, BL, blue). (**c**) Multivariate analysis comparing overall survival probabilities of BL patients (n = 34, GSE4475) when divided into 4 groups based on combined levels of STAT1 and STAT2. (**d**) Proportions of patients with no activated NK cells (estimated by CIBERSORT) when BL patients (n = 34, GSE4475) are divided into 4 groups based on combined levels of STAT1 and STAT2. **(e)** Comparison of overall survival probabilities of BL patients (n = 34, GSE4475) separated into three categories based on levels of activated NK cells in the tumor estimated by CIBERSORT, as none (0%, no activated NK), intermediate (Activated NK^Intermediate^), and high (Activated NK^High^). **(f-h)** Comparison of mRNA levels of MYC (**f**), STAT1 (**g**), and STAT2 (**h**) in mononuclear cells isolated from healthy human blood (n = 20) and from blood of T-ALL patients (n = 48, GSE62156). **(i-l)** CIBERSORT estimates of relative proportions of T cell blasts (**i**), and NK subsets, namely, total NK (**j**), resting NK (**k**) and activated NK (**l**), in peripheral blood mononuclear cells isolated from healthy individuals (n = 20) and T-ALL patients (n = 48, GSE62156). P-values for all survival analyses have been calculated using the log-rank test. P-values for all other analyses have been calculated using Mann-Whitney test. P-values: ns = not significant; *p < 0.05; **p < 0.01; ****p < 0.0001.

Next, we analyzed whether separation of BL patients into four groups based on their median expressions of both STAT1 and STAT2 can predict clinical prognosis. We observed that patients with the lowest levels of both STAT1 and STAT2 had the most unfavorable clinical outcome (Figure 6c). We observed that majority (85%, 6/7) of BL patients (n = 34) with lowest levels of both STAT1 and STAT2 have no activated NK cells (Figure 6d), as estimated by CIBERSORT. We found that higher levels of activated NK cells were associated with a significantly favorable clinical outcome and prolonged BL patient survival (Figure 6e). Hence, we conclude that low expression of both STAT1 and STAT2 (STAT1_Low_ STAT2_Low_) coincides both with maximum NK cell suppression (No activated NK, Figure 6d) and poor overall survival in BL patients (Figures 6c, 6e).

Finally, using CIBERSORT on bulk expression data, we compared the estimated fractions of NK cells in mononuclear cells isolated from blood of oncogene-addicted human T-lymphoma patients (n = 48, GSE62156) ^44,45^ and normal healthy individuals (n = 20) ^22^. Of note, MYC levels are significantly higher in T cell lymphoma patients as compared to healthy individuals (Figure 6f) because of chromosomal translocations that can potentially activate MYC ^44–50^. We observed a significant suppression of STAT1 and STAT2 levels (Figures 6g-h), and a reduction in total and activated NK cell subsets in human T cell lymphoma patients in comparison to their normal counterparts (Figures 6i-l). Collectively, our findings suggest a common paradigm of NK cell suppression in both human and mouse MYC-driven lymphomas, that occurs in part due to the direct repression of the STAT1/2-Type I IFN signaling cascade by the MYC oncogene.

### NK cells may represent a potential cell-based therapeutic against MYC-driven lymphomas

Our findings thus far suggested that subversion of NK cell-mediated immune surveillance is critical for MYC-driven lymphomagenesis. Next, we investigated whether NK cells can be developed as an immunotherapy against MYC-driven lymphomas by evaluating the effects of NK cells in MYC-driven lymphoma initiation, and recurrence post MYC inactivation.

First, we injected intravenously luciferase-labeled MYC-driven T cell lymphoblasts derived from an *SRα-tTA/tet-O-MYC* mouse (**Supplementary Figure 14**) into recipients with (NOD SCID) or without (NOD SCID IL-2R**γ**^-/-^, NSG) NK and other innate lymphoid cells. Lymphoma onset (day 5 for NOD SCID IL-2R**γ**^-/-^ and day 18 for NOD SCID, Figures 7a-b), and median morbidity (day 10 for NOD SCID IL-2R**γ**^-/-^ and day 20 for NOD SCID, Figure 7b) of mice were significantly delayed in NOD SCID transplant recipients with NK cells. T cell lymphomas eventually engrafted in the bone marrow of NOD SCID mice where NK cells are present (Figure 7a). We infer that while NK cell-mediated immune surveillance initially blocks MYC-driven lymphomagenesis, continued MYC expression eventually suppresses NK cells to establish lymphomas, as seen in primary *SRα-tTA/tet-O-MYC* mice.

**Figure 7:**
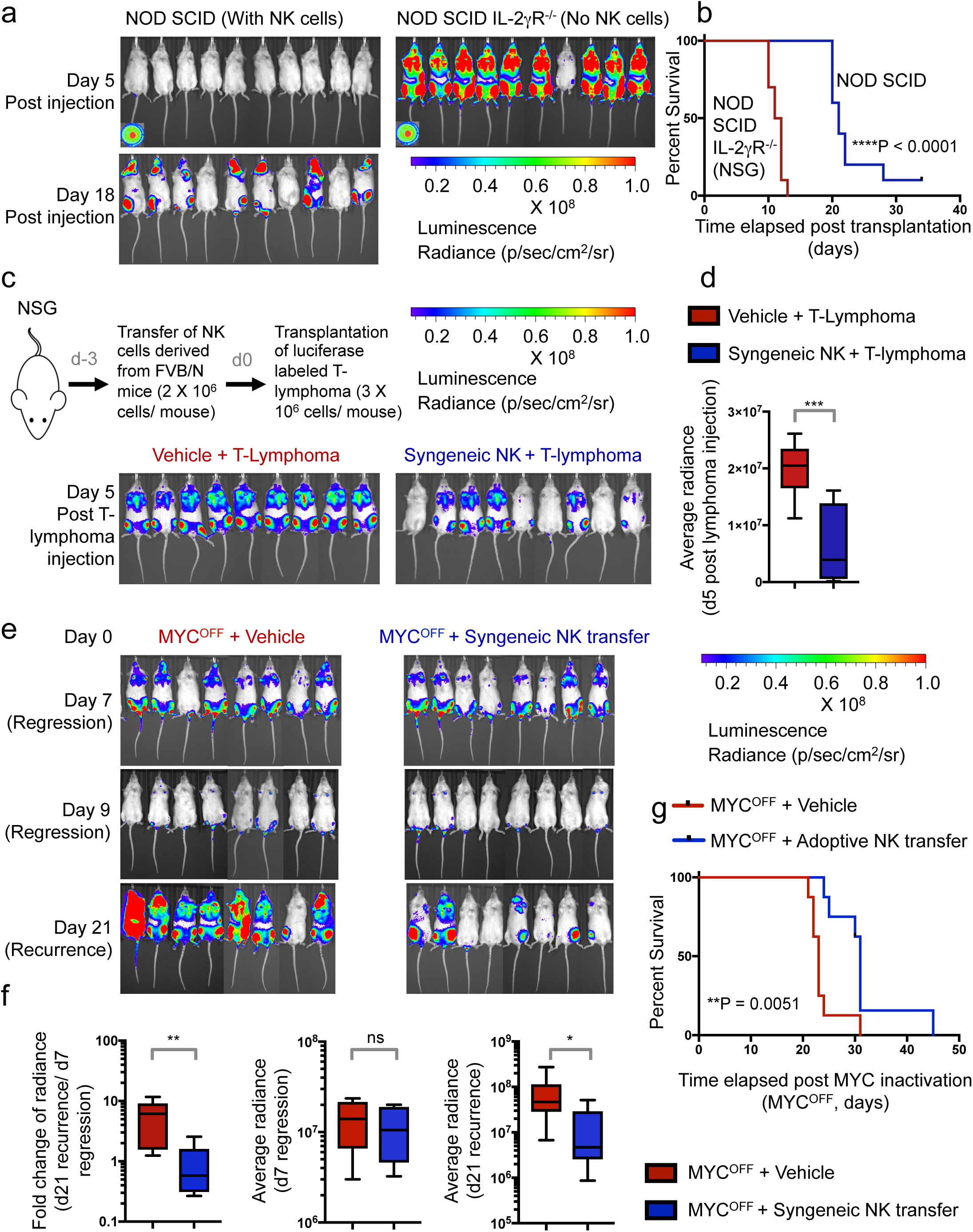
NK cells slow down growth and recurrence of MYC-driven lymphomas. **(a)** Bioluminescence imaging (BLI) of *NOD-SCID* (n = 10) and *NOD-SCIDIL2Rγ*^*-/-*^ (n = 10) mice transplanted with SR*α*-tTA MYC^ON^ mouse T-lymphoma cells post injection to monitor lymphoma engraftment and initiation. **(b)** Comparison of survival probabilities of *NOD-SCID* (n = 10) and *NOD-SCIDIL2Rγ*^*-/-*^ (NSG, n = 10) T-lymphoma transplant recipients. **(c)** BLI of NSG T cell lymphoma transplant recipients comparing lymphoma engraftment and initiation in the presence (n = 9) and absence (n = 9) of NK cells syngeneic to the tumor. (**d**) Quantification of changes in BLI signal of NSG transplant recipients that received vehicle or syngeneic NK cells before transplantation of T-lymphoma cells. **(e)** BLI of NSG T cell lymphoma transplant recipients comparing recurrence rates upon MYC inactivation in the absence (MYC^OFF^ + vehicle, n = 8) or presence of NK cells (MYC^OFF^ + Adoptive NK transfer, n = 8). **(f)** Quantification of changes in BLI signal (left) and absolute signals (middle and right) at the time of lymphoma regression (d7 MYC^OFF^) and lymphoma relapse (d21 MYC^OFF^). **(g)** Comparison of survival probabilities of NSG T-lymphoma transplant recipients that received either vehicle (n = 8) or adoptive NK transfer (n = 8) at the time of MYC inactivation (MYC^OFF^). P-values for all survival analyses have been calculated using the log-rank test. P-values for all other analyses have been calculated using Mann-Whitney test. P-values: ns = non significant; *p < 0.05; **p < 0.01; ****p < 0.0001.

Engraftment of T cell lymphoma at sites with NK cell-mediated surveillance such as the bone marrow (Figure 7a, BLI image day 18 post injection) suggested that NK cell-mediated allogeneic rejection is an unlikely explanation for delayed lymphoma engraftment in NOD SCID mice. However, we wanted to definitively rule out this possibility by transplanting syngeneic mature NK cells (CD3^-^ NKp46^+^) obtained from healthy normal mice (**Supplementary Figure 18**) into NSG mice three days before (d-3) T cell lymphoma transplantation (d0) (Figure 7c). We observed that T cell lymphoma initiation is significantly delayed in transplant recipients that received syngeneic NK cells in comparison to transplant recipients that received the vehicle control (Figures 7c-d). Despite delayed lymphoma initiation in transplant recipients that received NK cells, systemic lymphoma eventually develops consistent with our previous findings that, once established, MYC-driven lymphomas promote NK cell suppression.

MYC-driven lymphomas recur post MYC inactivation with 100% penetrance in immune-deficient hosts (NOD SCID IL-2R**γ**^-/-^) in comparison to immune-competent hosts where MYC inactivation sustains lymphoma regression ^13,51^. We investigated whether NK cells are sufficient to delay lymphoma recurrence post MYC inactivation. Syngeneic NK cells (CD3^-^ NKp46^+^) from normal mice when adoptively transferred into T cell lymphoma-bearing NOD SCID IL-2R**γ**^-/-^ recipients at the time of MYC inactivation delayed lymphoma recurrence (Figures 7e-f), and prolonged overall survival (Figure 7g) when compared to vehicle-treated controls. Thus, NK cells sustain lymphoma regression post MYC inactivation. Collectively, our results suggest that NK cells play an anti-tumorigenic role in MYC-driven lymphoid malignancies, hence making NK cell-based immune therapies as an attractive treatment strategy against MYC-driven lymphomas.

## Discussion

Mapping the perturbation in immune cell subsets during primary tumorigenesis is crucial for identifying immunotherapies that will most effectively treat specific cancers. We employed CyTOF to study the global immune cellular signature of overt MYC-driven T cell lymphomas before and after MYC inactivation (Figure 1, **Supplementary Figures 2-4**). We identified a novel mechanism by which oncogenic MYC oncogene directly suppresses NK cell-mediated immune surveillance. MYC-dependent suppression of NK cells was highly specific since other anti-tumorigenic immune subsets were not altered in numbers by both MYC activation and inactivation (Figure 1, **Supplementary Figures 2-4**). Notably, our results are consistent with a recent report where oncogenic MYC has been shown to promote NK cell exclusion in adenocarcinomas ^12^. Our studies provide a novel mechanism by which oncogenic MYC orchestrates such NK exclusion from lymphomas (Figures 3-5), illustrate human relevance (Figure 6), and demonstrate the therapeutic potential of NK cell adoptive transfers for treating MYC-driven lymphomas (Figure 7).

We found that mice bearing overt MYC-driven lymphomas have reduced number of NK cells because of a developmental blockade at the NK precursor (NKP) stage that prevents the production of more mature NKp46^+^ iNK cells (Figure 3). MYC inactivation significantly reversed this effect. Hence, we conclude MYC-driven lymphomas evade NK cell-mediated immune surveillance by blocking NK cell maturation, and infer that inhibiting MYC is essential for reversing this developmental blockade and eliciting an NK cell-mediated response against “MYC-addicted” tumors. Arrest in NK cell maturation has been demonstrated to be a critical mechanism of immune escape in cancers ^26,27^. Our findings demonstrate that the overexpression of oncogenic MYC in malignant lymphocytes orchestrates such arrest in NK cell development in the lymphoma microenvironment.

We show that MYC-driven lymphomas evade NK cell-mediated immune surveillance by transcriptionally repressing the STAT1/2-Type I IFN cytokine axis (Figures 5g-k, **Supplementary Figure 17**) required for NK cell maturation ^29–35^. Malignant lymphocytes overexpressing oncogenic MYC show suppressed production of Type I IFNs (Figures 5e-f, **Supplementary Figures 10, 15**). In turn, this blocks the maturation of NK cells in the lymphoma microenvironment. We also found that MYC-driven human lymphomas were associated with concomitant suppression of STAT1/2 and exclusion of NK cells (Figure 6).

We could restore NK cell production in spleens of lymphoma-bearing mice upon Type I IFN treatment (Figures 4f-j). Hence, MYC suppresses the STAT1/2-Type I IFN cascade and blocks NK cell-mediated immune surveillance (**Supplementary Figure 19**). We infer that administration of Type I IFNs can bypass the maturation defect of NK cells and restore NK cell-mediated immune surveillance of MYC-induced hematologic malignancies. In addition to their use as anti-viral agents, Type I IFNs can elicit anti-proliferative effects on cancer cells ^33,52^. Our current study explains why MYC-driven B cell lymphoma lines are sensitive to Type I IFN therapy ^53^, and illustrates new functional roles for Type I IFN therapy in lymphoid malignancies. Prior studies have shown that specific NK cell receptor-ligand interactions mediate the recognition of tumors by NK cells and can block MYC-driven lymphomagenesis ^54^. Expression levels of NKG2D (KLRK1) that were reduced *in vivo* in our primary autochthonous model bearing overt lymphoma (MYC^ON^), were restored to normal levels post MYC inactivation (MYC^OFF^) (Figure 4e), suggesting a possible suppression of NKG2DL on MYC-driven tumor cells. Consistent with a previous study, we observe a transcriptional repression of B2M in MYC-driven primary murine lymphomas (MYC^ON^) in comparison to healthy controls and MYC-inactivated (MYC^OFF^) lymphomas (Figure 4e) ^55^. Therefore, multiple mechanisms of MYC-mediated suppression of NK surveillance may be at play during lymphomagenesis.

Finally, our results may have implications for cancer immunotherapy ^56^. We observed that adoptively transferred NK cells significantly delayed overt MYC-driven lymphomagenesis, and contributed to sustained lymphoma regression post MYC inactivation (Figure 7), suggesting that MYC-driven lymphomas may be particularly sensitive to NK cell-based therapies. Most conventional chemotherapies used for lymphoid malignancies are immune suppressive. Therapies that are directed at restoring the immune response may be particularly effective against MYC-driven hematopoietic malignancies. Our findings underscore the importance of identifying MYC inhibitors ^57^ and combining these with NK cell-based therapies ^58^ to achieve maximum therapeutic benefit. We suggest that, as others and we investigate potential therapeutic candidates that target MYC, it will be essential to consider the role of NK cell-mediated immune surveillance in defining the modes of therapy; and to identify the characteristics, timing, and dosage of NK cells for maximum therapeutic benefit.

## Methods

### Cell lines and cell culture

Conditional MYC-driven mouse T cell lymphoma cell lines (**Supplementary Figure 14**) were derived from *SRα-tTA/tet-O-MYC* mice. c-MYC was inhibited in *SRα-tTA/tet-O-MYC* T-lymphoma cells by treating cell cultures with 0.02 µg/ml doxycycline (Sigma-Aldrich, T7660) for 4, 8, 24 and 48h. BL-like human P493-6 cells were kindly provided by Chi Van Dang, University of Pennsylvania. For c-MYC inactivation in P493-6 cells, the conditional p*myc*-tet construct was repressed with 0.1 µg/ml doxycycline (Sigma-Aldrich, T7660) for 24 h. Cell lines were confirmed to be negative for mycoplasma contamination and maintained in Roswell Park Memorial Institute 1640 medium (RPMI, Invitrogen) with GlutaMAX containing 10% fetal bovine serum, 100 IU/ml penicillin, 100 µg/ml streptomycin, 50 mM 2-mercaptoethanol at 37 °C in a humidified incubator with 5% CO_2_.

### Isolation and processing of immune cells from mice

We used young male mice (around 8-12 weeks of age). The following strains of mice were used: *SRα-tTA/tet-O-MYC* mice and the corresponding FVB/N wildtype strain. The *SRα-tTA/tet-O-MYC* males developed T cell lymphoma at approximately two-three months of age. Diseased mice were identified based on the presence of visible symptoms of T cell lymphoma development; namely, loss of weight and appetite, ruffled fur, shortness of breath and movement difficulties. Half the number of male littermates that developed T cell lymphoma were subjected to doxycycline treatment to inactivate MYC. For T cell lymphoma mice subject to MYC inactivation, one starter dose of doxycycline was given intraperitoneally, after which, the mice continuously received doxycycline for four days in their drinking water. Leukemic littermates from each cage were randomly and equally divided between MYC^ON/HIGH^ (untreated) or MYC^OFF/LOW^ (doxycycline-treated) groups. Spleens, blood and bone marrow were isolated from the three mice groups, namely, *SRα-tTA/tet-O-MYC* (MYC^ON/HIGH^), *SRα-tTA/tet-O-MYC* + Doxycycline (MYC^OFF/LOW^), and normal healthy (FVB/N) controls. Blood was drawn by cardiac puncture after euthanizing the mice, and directly subjected to erythrocyte lysis. Bone marrow was obtained by flushing the cavities of femur and tibia with PBS. Spleen and bone marrow were homogenized using a 19^1^/^2^G needle and filtered through a 70µm filter. After depletion of erythrocytes from spleen, blood and bone marrow using RBC lysis buffer (BD PharmLyse, BD Biosciences), washed cells were cryopreserved and utilized for CyTOF or flow cytometry. No blinding was done while executing the animal experiments.

### Mass cytometry (CyTOF Immunophenotyping)

This assay was performed in the Human Immune Monitoring Center at Stanford University. Mouse splenocytes were thawed in warm media, washed twice, resuspended in CyFACS buffer (PBS supplemented with 2% BSA, 2 mM EDTA, and 0.1% soium azide). Viable cells were then counted by Vicell. Cells were added to a V-bottom microtiter plate at 1 million viable cells/well and washed once by pelleting and resuspension in fresh CyFACS buffer. The cells were stained for 60 min on ice with 50uL of the following antibody-polymer conjugate cocktail (**Supplementary Table 1**). All antibodies were custom-ordered metal-conjugates from Fluidigm (**Supplementary Table 1**). The cells were washed twice; by pelleting and resuspension with 500 uL FACS buffer. The cells were resuspended in 100uL PBS buffer containing 1.7 ug/mL Live-Dead (DOTA-maleimide (Macrocyclics) containing natural-abundance indium). The cells were washed twice by pelleting and resuspension with 500uL CyFACS buffer. The cells were resuspended in 100uL 2% PFA in PBS and placed at 4°C overnight. The next day, the cells were pelleted and washed by resuspension in fresh PBS. The cells were resuspended in 100uL eBiosciences permeabilization buffer (1x in PBS) and placed on ice for 45 min before washing twice with 500uL CyFACS buffer. The cells were resuspended in 100uL iridium-containing DNA intercalator (1:2000 dilution in PBS; DVS Sciences) and incubated at room temperature for 20 min. The cells were washed once in CyFACS buffer, then three times in 500uL MilliQ water. The cells were diluted to a concentration of ∼800,000 cells/mL in MilliQ water containing 0.1x EQ normalization beads (Fluidigm) before injection into the CyTOF (Fluidigm). Data analysis was performed using FlowJo v10 by gating on intact cells based on the Iridium vs Ce140 (bead), then both Iridium isotopes from the intercalator, then on singlets by Ir191 vs cell length, then on live cells (Indium-LiveDead minus population), followed by cell subset-specific gating (**Supplementary Figure 20**). The data was analyzed using FlowJo X 10.0.7r2 and viSNE (Cytobank Inc.).

### Phospho-CyTOF

The harvested cells were suspended in 16% PFA and incubated for 10 minutes at room temperature. After washing with CyFACS buffer (1x CyPBS with 0.1% BSA, and 0.05% Na Azide; made in Milli-Q water), surface staining was performed by incubating the cells with the surface-staining antibody cocktail (**Supplementary Table 2**) at room temperature for 30 minutes. The cells were then washed with CyFACS buffer and permeabilized before carrying out intracellular staining. The cells were then incubated with the intracellular antibody cocktail (**Supplementary Table 2**) at room temperature for 30 minutes. After washing with CyPBS, the cells were incubated with Ir-intercalator solution on ice for 20 minutes. The samples were then washed with Milli-Q water and acquired on the CyTOF instrument using standard instrument set up procedures. The data was analyzed using FlowJo X 10.0.7r2.

### Flow cytometry

All antibodies for flow cytometry measurements as well as their respective isotype controls are indicated in **Supplementary Table 3**. Surface staining antibodies that were PE conjugated were used at a dilution of 1:10 and the ones conjugated with FITC and APC at a dilution of 1:5. Multicolor flow cytometry staining was performed according to standard procedures. Cells were stained with Propidium Iodide (PI, Sigma-Aldrich) to gate out dead cells. Labeled cells were analyzed on a FACSCalibur (Becton Dickinson). The data was analyzed using FlowJo X 10.0.7r2. Gating strategy is provided in **Supplementary Figure 20**.

### Calculation of absolute immune cell counts

Viable cells were counted using a Vicell before subjecting the single cell suspensions of spleen, blood and bone marrow to CyTOF or flow cytometry, respectively. Using the percentages of the immune populations calculated by CyTOF and flow cytometry on the same day, absolute viable cell numbers of the different immune subsets were computed.

### RNA sequencing

A part of the splenic single cell suspensions from normal (FVB/N, n =3), SR*α-*tTA MYC^ON^ (n = 3) and SR*α*-tTA MYC^OFF^ (n = 3) mice subjected to CyTOF were used to generate RNA. RNA sequencing was performed at the Beijing Genomics Institute (BGI) using the BGISEQ-500 instrument. The experimental pipeline is described below:

#### Library Preparation

Oligo (dT) magnetic beads were used to select mRNA with polyA tail, or hybridize the rRNA with DNA probe and digest the DNA/RNA hybrid strand, followed by DNase I reaction to remove DNA probe. After obtaining the target RNA, it was fragmented and reverse transcribed to double-strand cDNA (dscDNA) by N6 random primer. The dscDNA was end repaired with phosphate at 5’ end and stickiness ‘A’ at 3’ end. The adaptor with stickiness ‘T’ at 3’ end was ligated to the dscDNA. Two specific primers were used to amplify the ligation product. The PCR product was then denatured and the single strand DNA was cyclized by splint oligo and DNA ligase. Sequencing was then performed on the prepared library.

#### Bioinformatics Pipeline

Primary sequencing data (raw reads) were subjected to quality control (QC). After QC, raw reads were filtered into clean reads before alignment to the reference sequences. Once alignment passed QC, we proceeded with downstream analysis including gene expression and deep analysis based on gene expression (PCA/correlation/screening differentially expressed genes).

#### Data Filtering

Data was filtered reads with adaptors, reads with greater than 10% unknown bases and low quality reads.

#### Reads Mapping

Bowtie 2 was used to map clean reads to reference gene, and HISAT was used to map reads to reference genome (mm8).

#### Gene Quantification

RSEM was used to compute Maximum likelihood abundance estimates using the Expectation Maximization (EM) algorithm. FPKM method was used to calculate the expression levels of individual genes.

RNA sequencing data are provided in Gene Expression Omnibus (GSE106078).

### CIBERSORT

Human CIBERSORT was carried out at https://cibersort.stanford.edu/ using LM22 reference matrix of gene expression in human immune subsets *(19)*. For CIBERSORT of splenic immune compositions in normal (FVB/N), SR*α-*tTA MYC^ON^ and SR*α-*tTA MYC^OFF^ mice, a reference matrix was generated using ImmGen (https://www.immgen.org/, GSE15907). CIBERSORT was carried out on CyTOF-matched normal (n =3), SR*α-*tTA MYC^ON^ (n = 3) and SR*α-*tTA MYC^OFF^ (n = 3) samples.

### Transplantation of MYC-addicted T cell lymphoma and bioluminescence imaging

A MYC-addicted T cell lymphoma cell line derived from a male *SRα-tTA/tet-O-MYC* mouse (**Supplementary Figure 14**) was labeled with luciferase. 3 × 10^6^ luciferase-labeled T cell lymphoma cells were then injected intravenously into two groups of immune-deficient hosts: NOD SCID (The Jackson Laboratory) and NOD-SCIDIL-2R**γ**^-/-^ (NSG). Male transplant recipients used were between 4-5 weeks of age. Leukemic engraftment and progression were tracked by bioluminescence imaging (BLI) of the transplant recipients twice every week using an *in vivo* IVIS 100 bioluminescence/optical imaging system (Xenogen). D-Luciferin (Promega) dissolved in PBS was injected intraperitoneally at a dose of 2.5 mg per mouse 15 min before measuring the luminescence signal. General anesthesia was induced with 3% isoflurane and continued during the procedure with 2% isoflurane introduced through a nose cone. No blinding was done while executing the animal experiments. All mouse experiments were approved by the Administrative Panel on Laboratory Animal Care (APLAC) at Stanford University, and were carried out in accordance with institutional and national guidelines.

### Adoptive transfer of NK cells into T cell lymphoma-bearing recipient mice

3 × 10^6^ luciferase-labeled T cell lymphoma cells were injected intravenously into 4-8 week old NSG mice (n=19). Post verification of T cell lymphoma engraftment by BLI, all mice were treated with doxycycline to inactivate MYC. 10 of the 19 mice were subjected to adoptive transfer of NK cells isolated from spleens of 4-8 week old FVB/N mice by magnetic activated cell sorting (MACS, NK Cell Isolation Kit II, Miltenyi Biotech). Flow cytometry was performed to confirm the purity of the isolated NK cells. 1 × 10^6^ MACS purified NK cells were injected per mouse into NSG transplant recipients bearing regressed T cell lymphoma (on doxycycline, MYC^OFF^).

### Type I IFN treatment of primary *SRα-tTA/tet-O-MYC* mice

20,000U of Type I IFN*α* (R&D systems) was administered intraperitoneally to SR*α-tTA/tet-O-MYC* mice bearing overt lymphoma for 3 days. On day 3, spleens from treated mice were collected, and subjected to flow cytometry for NK and T cells. PBS-treated mice were used as controls. Absolute cell counts were obtained as described previously.

### Immunoblotting

Cells were lysed in CelLytic buffer (Sigma) supplemented with 1% protease inhibitor ‘cocktail’ (Pierce). Protein samples were subsequently separated by electrophoresis through NuPAGE (Invitrogen) 4–12% Bis-Tris gradient gels and were transferred to PVDF membranes (Immobilion; Millipore). For the detection of mouse and human proteins by immunoblot analysis, we used primary antibodies together with the WesternBreeze immunodetection system (Invitrogen). All antibodies used for immunoblotting and their corresponding dilutions are given

### Retrovirus production and transduction

Transfection of retroviral constructs (**Supplementary Table 5**) was performed using Lipofectamine 2000 (Invitrogen) with Opti-MEM media (Invitrogen). Retroviral supernatants for infection of cells were generated after co-transfection of Phoenix cells containing the plasmids pHIT60 (gag-pol) and pHIT123 (ecotropic envelope) (provided by G.P. Nolan), with the retroviral constructs. Cells were cultured in high-glucose DMEM (Invitrogen) with GlutaMAX containing 10% FBS, 100 IU/ml penicillin, 100 µg/ml streptomycin, 25 mM HEPES, pH 7.2, 1 mM sodium pyruvate and 0.1 mM nonessential amino acids. Serum-free medium was replaced after 16 h with growth medium containing 10 mM sodium butyrate. After 8 h of incubation, the medium was changed back to growth medium without sodium butyrate. 24 h later, we harvested the viral supernatants, passed them through a 0.45-µm filter, and centrifuged them twice at 2,000*g* for 90 min at 32 °C on 50 µg/ml RetroNectin-coated non–tissue-culture-treated six-well plates. Pre-B cells (2 × 10^6^ to 3 × 10^6^ cells per well) were transduced by centrifugation at 600*g* for 30 min and were maintained overnight at 37 °C with 5% CO_2_ before transfer into culture flasks.

### Lentivirus production and transduction

Lentiviral (**Supplementary Table 5**) transfections were performed using Lipofectamine 2000 (Invitrogen) with Opti-MEM media (Invitrogen). We produced lentiviral supernatants to infect murine cells by cotransfecting HEK 293FT cells with the plasmids psPAX2 (gag-pol; Didier Trono, Addgene 12260) and pMD2.G (VSV-G envelope; Didier Trono, Addgene 12259). Cells were cultivated in high-glucose DMEM (Invitrogen) with GlutaMAX as described above. Culture supernatants were concentrated with Lenti-X™ Concentrator (Clontech) according to the manufactures recommendation. The concentrated viruses were incubated with the cells for 50 µg/ml RetroNectin-coated non-tissue culture treated 6-well plates.

### Quantitative RT-PCR

Quantitative real-time PCR carried out with the SYBRGreenER mix from Invitrogen according to standard PCR conditions and an ABI7900HT real-time PCR system (Applied Biosystems). Primers for quantitative RT-PCR are in **Supplementary Table 6**.

### Immunohistochemistry

Spleens and thymuses obtained from experimental mice were immersed in 20 ml of formalin (VWR) for 24h and transferred to PBS. Paraffin embedding was carried out using standard procedures on a Tissue-TEK VIP processor (Miles Scientific), and 4µm sections were mounted on Apex superior adhesive slides (Leica Microsystems) and stained on a Ventana BenchMark automated IHC stainer). Immunohistochemistry sections were mounted with mounting medium, antifade reagent (Pro-Long Gold; Invitrogen) was applied, and coverslips were sealed before acquisition of images at 25°C on a Zeiss Axiovert 200M inverted confocal microscope with a 40 Plan Neofluor objective using IP Lab 4.0 software (Scanalytics).

### ChIP-sequencing analysis

Raw ChIP-sequencing files were downloaded from Gene Expression Omnibus (GEO). The FASTQ Groomer in Galaxy Software (https://usegalaxy.org/) was used to generate files that were mapped to RefSeq database for mouse genes (mm9). The resulting bigWig files processed from Galaxy were visualized in Integrated Genome Viewer (IGV_2.3.72).

### Statistical analysis

CyTOF, flow cytometry and CIBERSORT results are shown as box plots and comparisons between any two groups were made using the Mann-Whitney U test (not assuming normal distributions) with Graphpad Prism software. All remaining data are presented as mean ± s.d. The comparisons for the mean values between two sample groups were made using the two-tailed Student’s *t*-test after calculating variances for each group with Graphpad Prism software. For experiments involving transplantation of cells into mice, the minimal number of mice in each group was calculated through the use of the ‘cpower’ function in the R/Hmisc package (R Development Core Team 2009; http://www.r-project.org). The Kaplan-Meier method was used for estimation of overall survival and relapse-free survival. For all survival analyses, the log-rank test was used to compare groups of transplanted mice or human patients. The R package ‘survival’ version 2.35-8 was used for the survival analysis. All p-values are two-tailed. The level of significance was set at *P* < 0.05. P-values: ns = non-significant; *p < 0.05; **p < 0.01; ***p < 0.001; ****p < 0.0001.

### Institutional Approval

All mouse experiments were approved by the Administrative Panel on Laboratory Animal Care (APLAC) at Stanford University, and were carried out in accordance with institutional and national guidelines.

## Author Contributions

S.S. and D.W.F. conceived the study. S.S. developed the experimental methodology and conducted most of the experiments. L.D.H., D.F.L., A.D., C.H., A.M., and M.L. performed experiments. R.D. helped with analysis and interpretation of RNA sequencing results. S.S. wrote the manuscript. D.W.F. and H.T.M. provided helpful scientific input on the manuscript. H.T.M., and D.W.F. helped in editing the manuscript. H.T.M and D.W.F. provided administrative, technical, and material support. D.W.F. supervised the study.

## Acknowledgements

We thank M. D. Leipold, R. Fernandez and Y. Rosenberg-Hasson at the Human Immune Monitoring Center, Stanford University for Luminex, CyTOF and Phospho-CyTOF acquisition. We also thank P. Chu in the Department of Pathology at Stanford University for carrying out tissue sectioning; H. Dai for technical help; A. Gentles for providing us access to PRECOG data sets; and M. Eilers for gifting us lentiviral overexpression plasmids for the study. We thank M. Müschen and M. Eilers for providing their expert scientific guidance and suggestions. This work is supported by the following grants: NIH/NCI CA170378PQ2, 1U01CA188383-01/S1, and Emerson Collective Cancer Research Fund to D. Felsher, and the Special Fellow Award from The Leukemia and Lymphoma Society (LLS 3366-17) and American Society of Hematology (ASH) Scholar Award to S. Swaminathan. D. F. Liefwalker is a recipient of the Burroughs Wellcome Fund Postdoctoral Enrichment Award.

## Conflicts of Interest

The authors declare no potential conflicts of interest.

