## Supplemental Figures and Tables for "MYC Functions as a Switch for Natural Killer Cell-Mediated Immune Surveillance of Lymphoid Malignancies"

### Inventory

- Supplementary Figures S1-S20
- Supplementary Tables S1-S6

**Supplementary Figure 1:** *MYC-driven lymphomas exhibit disrupted splenic architectures*

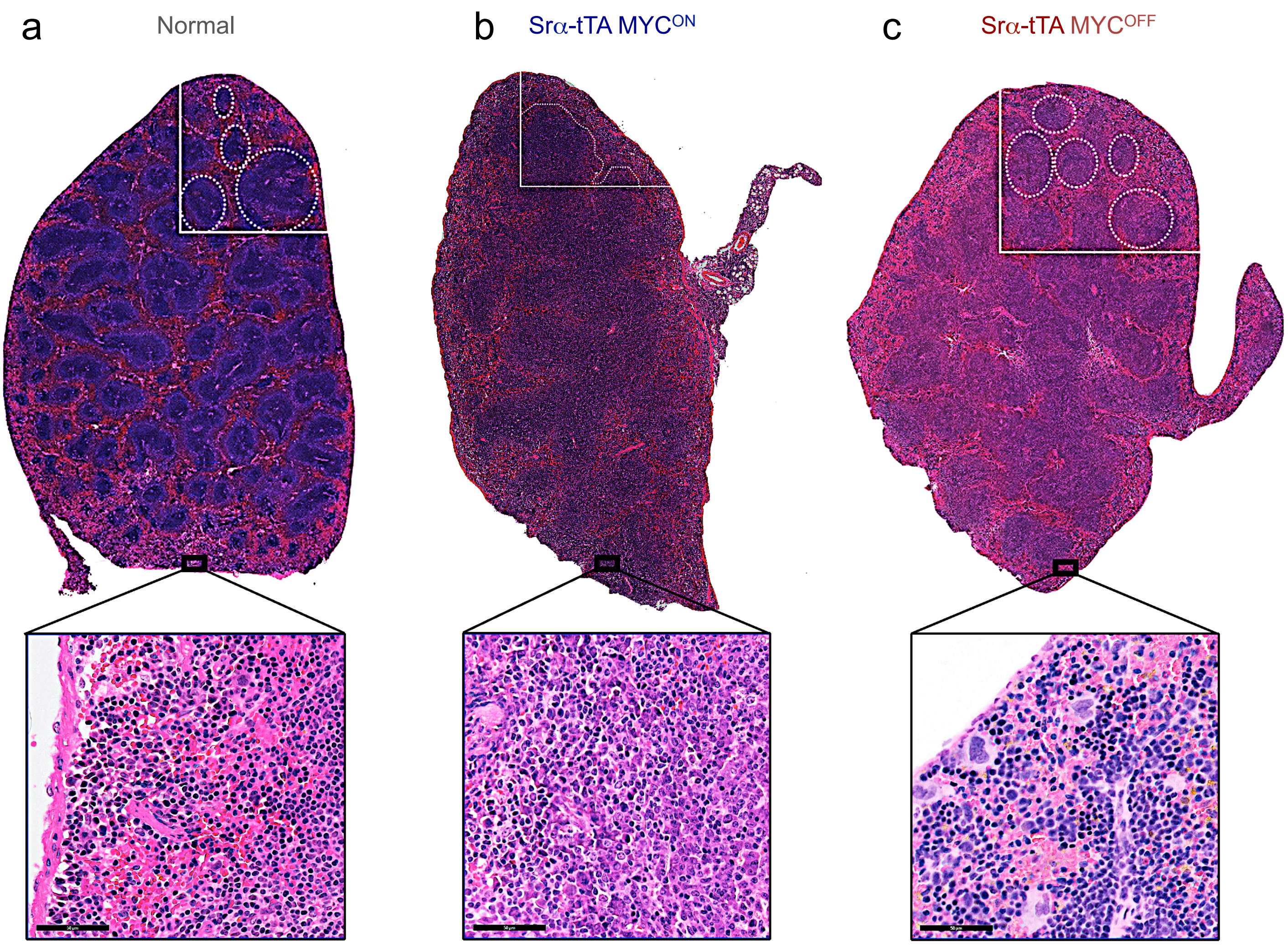


**Supplementary Figure 1: MYC-driven lymphomas exhibit disrupted splenic architectures.** Hematoxylin and Eosin (H&E) staining of spleens isolated from age and sex matched healthy normal (n = 3, **a**), Srα-tTA $MYC^{ON}$ (n = 3, **b**) and Srα-tTA $MYC^{OFF}$ (doxycycline 96 h, n = 3, **c**) mice. White dotted circles depict an intact germinal center in healthy and Srα-tTA $MYC^{OFF}$ mice. Magnification (top) = 4X, Inset (bottom): Magnification = 20X, Scale bars = 50 μm.

**Supplementary Figure 2:** *Oncogenic MYC perturbs the relative compositions of splenic immune subsets*

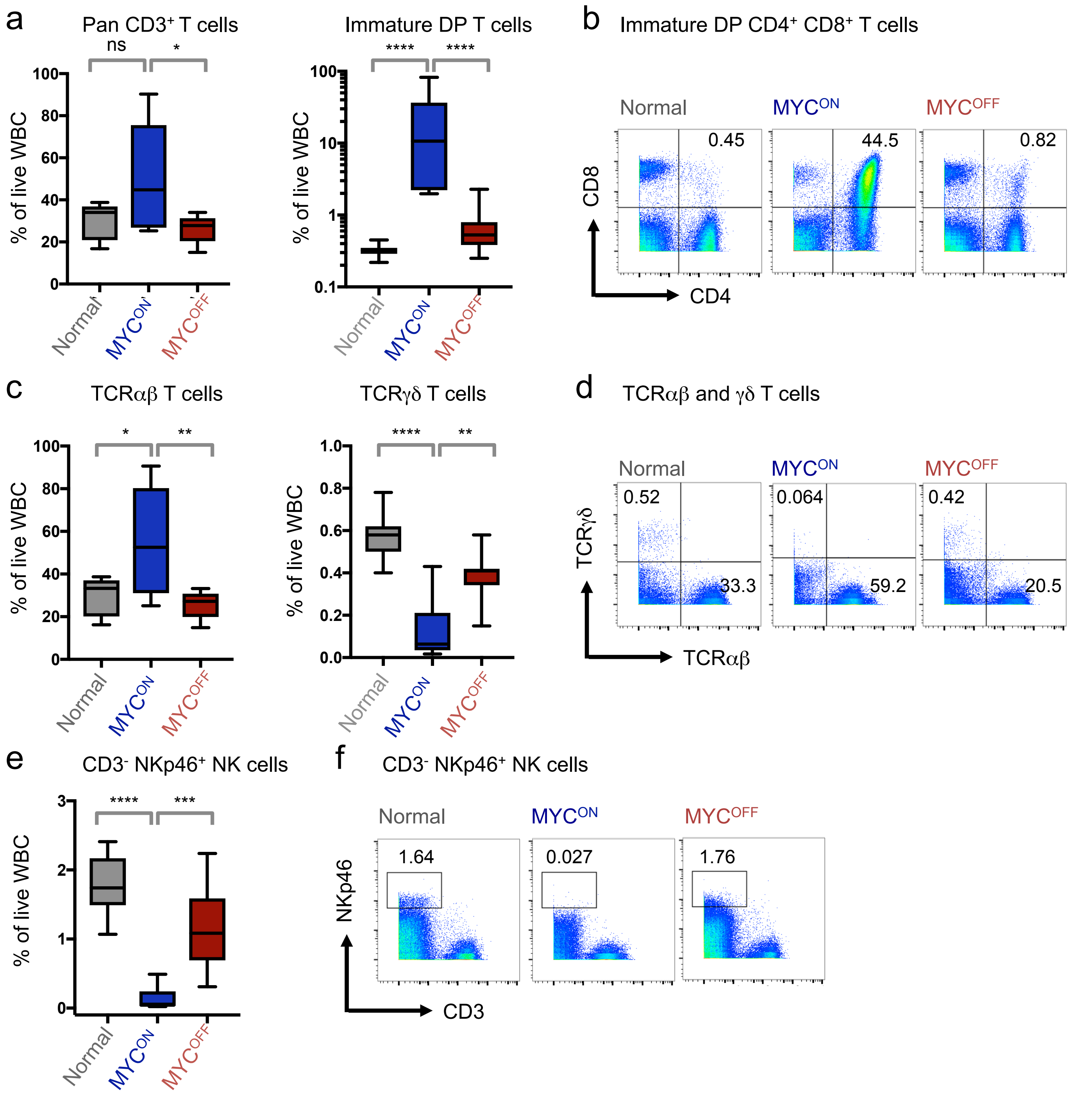


**Supplementary Figure 2: Oncogenic MYC perturbs the relative compositions of splenic immune subsets.** (**a**) Quantification of CD3⁺ T and CD4⁺ CD8⁺ immature DP T cell percentages in normal (n = 11), SRα-tTA MYC$^{ON}$ (n = 9), and SRα-tTA MYC$^{OFF}$ (doxycycline 96 h, n = 10) mice by CyTOF. (**b**) CyTOF dot plots showing percentages of DP T cells in one representative spleen from normal, SRα-tTA MYC$^{ON}$, and SRα-tTA MYC$^{OFF}$ mice. (**c**) Quantification of TCRαβ⁺ and TCRγδ⁺ T cell percentages in normal, SRα-tTA MYC$^{ON}$, and SRα-tTA MYC$^{OFF}$ mice. (**d**) CyTOF dot plots showing percentages of TCRαβ and TCRγδ T cells in one representative spleen from normal, SRα-tTA MYC$^{ON}$, and SRα-tTA MYC$^{OFF}$ mice. (**e**) Quantification of NK cell percentages in normal, SRα-tTA MYC$^{ON}$, and SRα-tTA MYC$^{OFF}$ mice. (**f**) CyTOF dot plots showing NK cell percentages in one representative spleen from normal, SRα-tTA MYC$^{ON}$, and SRα-tTA MYC$^{OFF}$ mice. All populations have been gated on live intact singlets. P-values were calculated using the Mann-Whitney test (ns = not significant, *$p < 0.05$, **$p < 0.01$, ***$p < 0.001$, ****$p < 0.0001$).

**Supplementary Figure 3:** *Oncogenic MYC perturbs the relative compositions of splenic immune subsets*

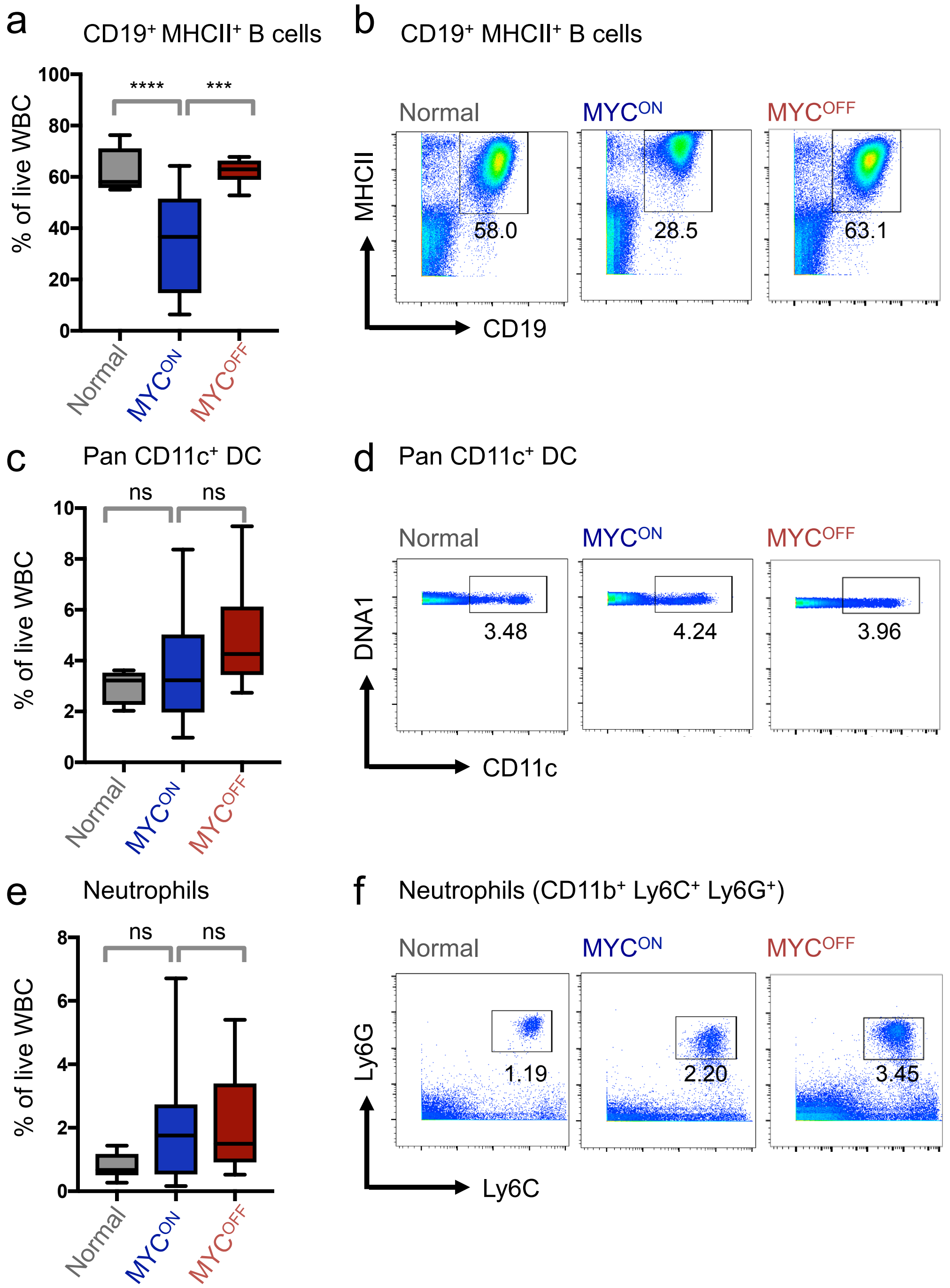


**Supplementary Figure 3: Oncogenic MYC perturbs the relative compositions of splenic immune subsets.** (**a**) Quantification of CD11c$^+$ DCs in normal (n = 11), SRα-tTA MYC$^{ON}$ (n = 9), and SRα-tTA MYC$^{OFF}$ (doxycycline 96 h, n = 10) mice by CyTOF. (**b**) CyTOF dot plots showing percentages of DCs in one representative spleen from normal, SRα-tTA MYC$^{ON}$, and SRα-tTA MYC$^{OFF}$ mice. (**c**) Quantification of neutrophil percentages in normal, SRα-tTA MYC$^{ON}$, and SRα-tTA MYC$^{OFF}$ mice. (**d**) CyTOF dot plots showing percentages of neutrophils in one representative spleen from normal, SRα-tTA MYC$^{ON}$, and SRα-tTA MYC$^{OFF}$ mice. (**e**) Quantification of B cell percentages in normal, SRα-tTA MYC$^{ON}$, and SRα-tTA MYC$^{OFF}$ mice. (**f**) CyTOF dot plots showing B cell percentages in one representative spleen from normal, SRα-tTA MYC$^{ON}$, and SRα-tTA MYC$^{OFF}$ mice. All populations have been gated on live intact singlets. P-values were calculated using the Mann-Whitney test (ns = not significant *$p < 0.05$, **$p < 0.01$, ***$p < 0.001$, ****$p < 0.0001$).

**Supplementary Figure 4:** *Changes in absolute numbers of splenic immune subsets during MYC-driven lymphomagenesis*

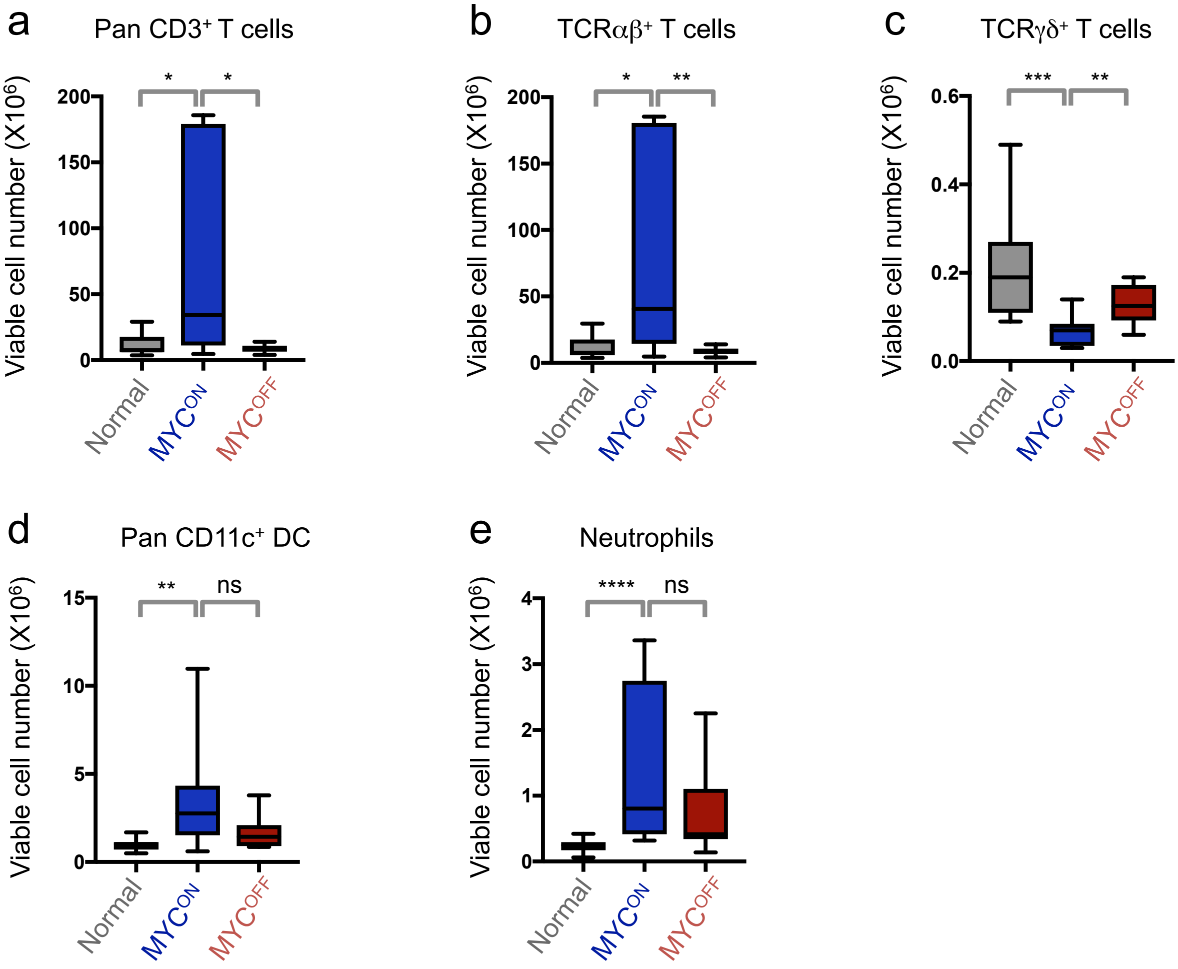


**Supplementary Figure 4: Changes in absolute numbers of splenic immune subsets during MYC-driven lymphomagenesis.** Quantification of absolute cell numbers of immune compartments in spleens of normal (n = 11), SRα-tTA $MYC^{ON}$ (n = 9), and SRα-tTA $MYC^{OFF}$ (doxycycline 96 h, n = 10) mice subjected to mass cytometry. P-values have been calculated using the Mann-Whitney test (ns = not significant, *p < 0.05, **p < 0.01, ***p < 0.001, ****p < 0.0001).

**Supplementary Figure 5:** *Alterations of splenic NK compositions in MYC-driven T cell lymphoma verified by flow cytometry*

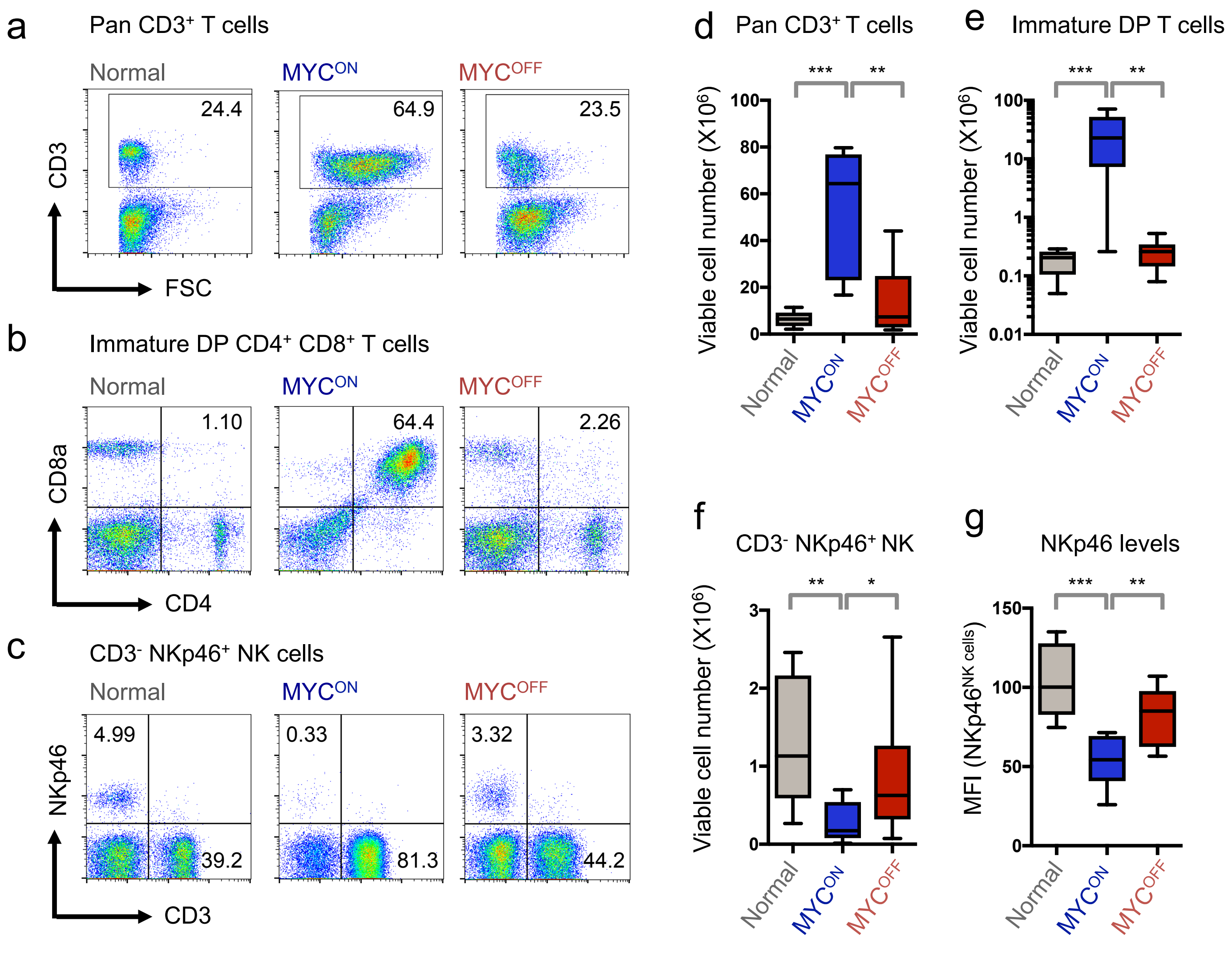


**Supplementary Figure 5: Alterations of splenic NK compositions in MYC-driven T cell lymphoma.** (**a-c**) Representative flow cytometry plots depicting splenic pan $CD3^+$ T cell (a), $CD4^+$ $CD8^+$ DP T cells (b), and $CD3^-$ $NKp46^+$ NK cell (c), distributions in normal (n = 8), SRα-tTA $MYC^{ON}$ (n = 8), and SRα-tTA $MYC^{OFF}$ (doxycycline 96 h, n = 8) mice. (**d-f**) Quantification of numbers of pan $CD3^+$ T cells (d), and immature DP T cell (e), and $CD3^-$ $NKp46^+$ NK cells (f) in normal (n = 8), SRα-tTA $MYC^{ON}$ (n = 8), and SRα-tTA $MYC^{OFF}$ (n = 8) spleens by flow cytometry. (**g**) MFI of surface NKp46 in splenic NK cells from normal (n = 8), SRα-tTA $MYC^{ON}$ (n = 8), and SRα-tTA $MYC^{OFF}$ (n = 8) mice. All populations are depicted as percentages of live cells ($FSC^+$ $PI^-$). P-values have been calculated using the Mann-Whitney test (ns = not significant, *$p < 0.05$, **$p < 0.01$, ***$p < 0.001$).

**Supplementary Figure 6:** *CIBERSORT measuring changes in splenic NK composition during murine MYC-driven T cell lymphomagenesis*

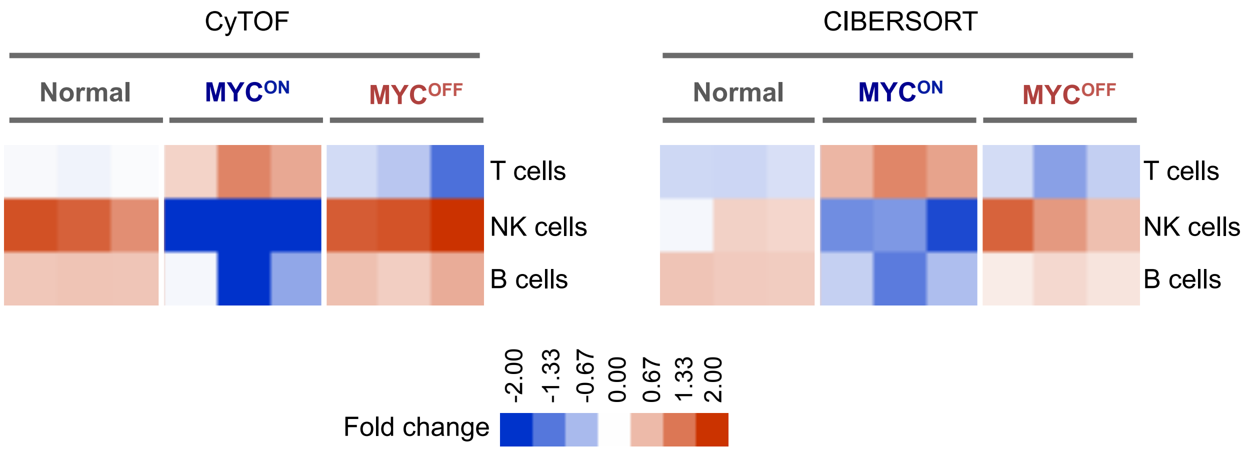


**Supplementary Figure 6: CIBERSORT measuring changes in splenic NK composition during murine MYC-driven T cell lymphomagenesis.** Heatmap comparing fold changes in T, B and NK immune subsets in normal (n = 3), SRα-tTA MYC$^{ON}$ (n = 3), and SRα-tTA MYC$^{OFF}$ (n = 3) mice by CyTOF (left) and CIBERSORT (right).

**Supplementary Figure 7:** *Suppression of bone marrow NK cells during MYC-driven lymphomagenesis*

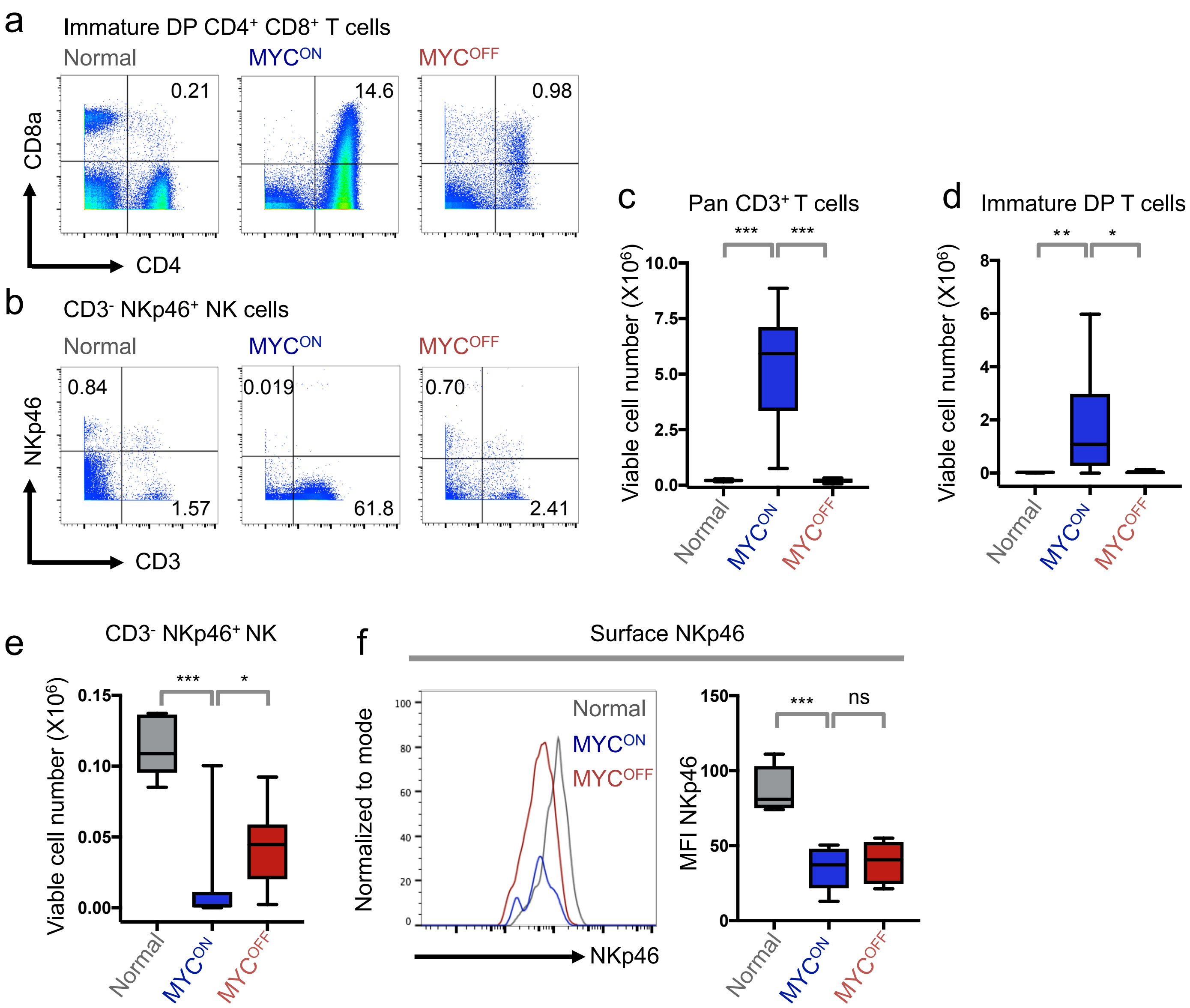


**Supplementary Figure 7: Suppression of bone marrow NK cells during MYC-driven lymphomagenesis.** (**a-b**) Representative CyTOF plots depicting T and NK cell distributions in normal (n = 7), SRα-tTA MYC$^{ON}$ (n = 9), and SRα-tTA MYC$^{OFF}$ (doxycycline 96 h, n = 7) bone marrows. (**c-e**) Quantification of absolute numbers of pan CD3$^{+}$ T cells (c), immature DP T cells (d), and NK cells (e) in bone marrows of normal, MYC$^{ON}$, and MYC$^{OFF}$ mice. (**f**) Representative MFI of surface NKp46 in bone marrow NK cells from normal, MYC$^{ON}$, and MYC$^{OFF}$ mice. All populations are depicted as percentages of live intact singlets. P-values have been calculated using the Mann-Whitney test (ns = not significant, *$p < 0.05$, **$p < 0.01$, ***$p < 0.001$).

**Supplementary Figure 8:** *Increased cell death is not responsible for reduction of NK cells in MYC-driven lymphomas*

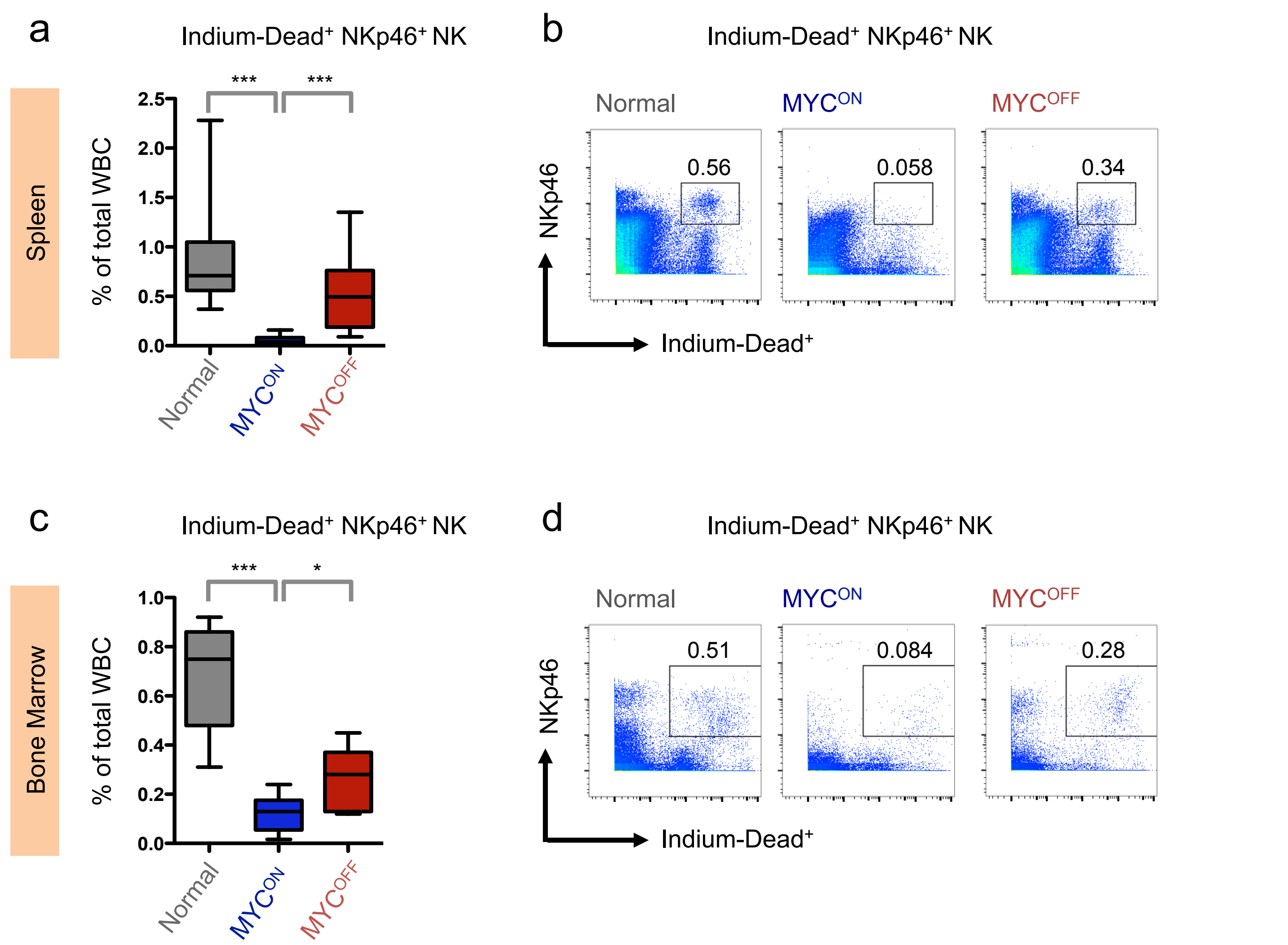


**Supplementary Figure 8: Increased cell death is not responsible for reduction of NK cells in MYC-driven lymphomas.** (**a-b**) Quantification of percentages (a), and representative plots (b) of dead NK cells (Indium-Dead$^+$ NKp46$^+$) in normal (n = 11), SRα-tTA MYC$^{ON}$ (n = 9), and SRα-tTA MYC$^{OFF}$ (doxycycline 96 h, n = 10) spleens measured by CyTOF. (**c-d**) Quantification of percentages (c), and representative plots (d) of dead NK cells (Indium-Dead$^+$ NKp46$^+$) in normal (n = 7), SRα-tTA MYC$^{ON}$ (n = 9), and SRα-tTA MYC$^{OFF}$ (doxycycline 96 h, n = 7) bone marrows measured by CyTOF.

**Supplementary Figure 9:** *Transcriptional repression of JAK-STAT signature during primary MYC-driven T cell lymphomagenesis*

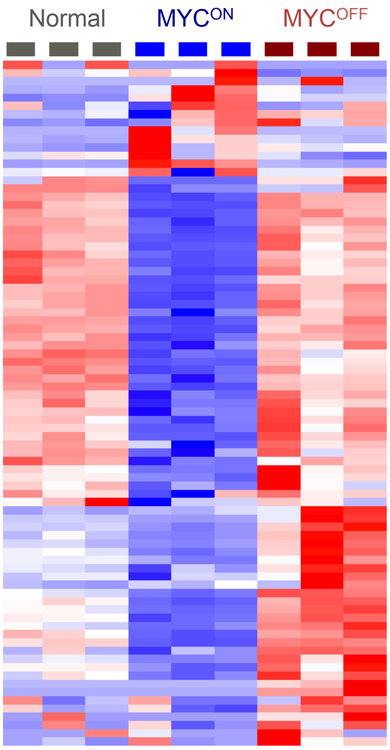


**Supplementary Figure 9: Transcriptional repression of JAK-STAT signature during primary MYC-driven T cell lymphomagenesis.** Heatmap showing suppression of JAK-STAT signaling in spleens of SRα-tTA MYC$^{ON}$ (n = 3) mice when compared to normal (n =3), and SRα-tTA MYC$^{OFF}$ (n = 3) mice.

**Supplementary Figure 10:** *Transcriptional repression of Type I IFN signaling during murine MYC-driven B cell lymphomagenesis*

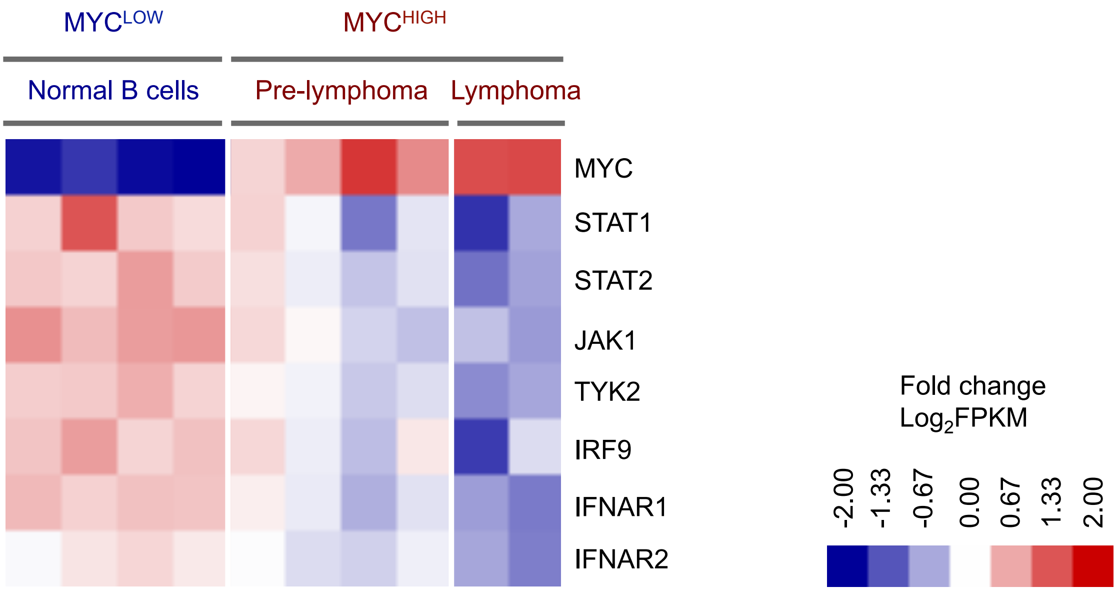


**Supplementary Figure 10: Transcriptional repression of Type I IFN signaling during murine MYC-driven B cell lymphomagenesis.** Heat map comparing transcriptional changes in Type I IFN signaling genes in normal B cells from healthy mice ($MYC^{LOW}$, n = 4), pre-lymphomagenic B cells from Eμ-MYC mice (pre-lymphoma, $MYC^{HIGH}$, n = 4), and in full blown primary B cell lymphomas from Eμ-MYC mice ($MYC^{HIGH}$, n = 2) using RNA sequencing (GSE 51008).

**Supplementary Figure 11:** *Signaling and surface receptor changes after MYC inactivation in human BL model*

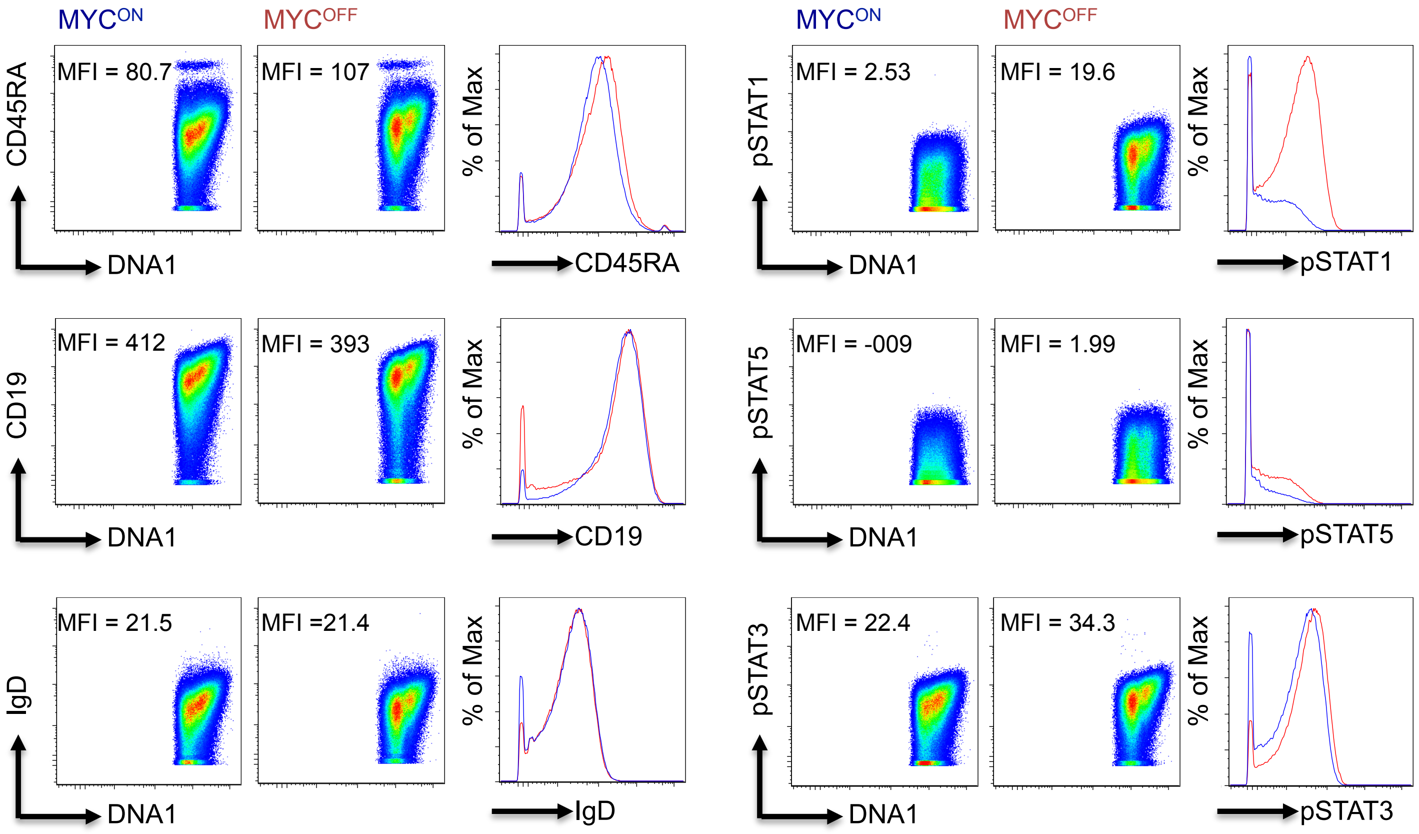


**Supplementary Figure 11: Signaling and surface receptor changes after MYC inactivation in human BL model.** Dot plots of Phospho-CyTOF depicting the global changes in B cell specific surface receptors and cytokine JAK-STAT signaling pathways after turning MYC off for 24h in P493-6 cells.

**Supplementary Figure 12:** *High MYC levels in B cell lymphomas block transcription of STAT1 and STAT2*

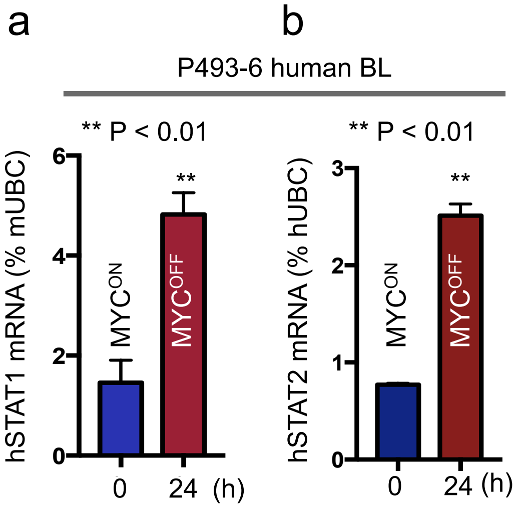


**Supplementary Figure 12: High MYC levels in B cell lymphomas block transcription of STAT1 and STAT2. (a-b)** Quantitative real time PCR for hSTAT1 (a) and, hSTAT2 (b) in P493-6 human BL cell line before and after MYC inactivation by doxycycline for 24 hours (n = 3, mean ± s.d). P-values have been calculated using the Student's t-test (*p < 0.05, **p < 0.01, ***p < 0.001).

**Supplementary Figure 13:** *MYC inactivation in BL changes the secreted cytokine profile*

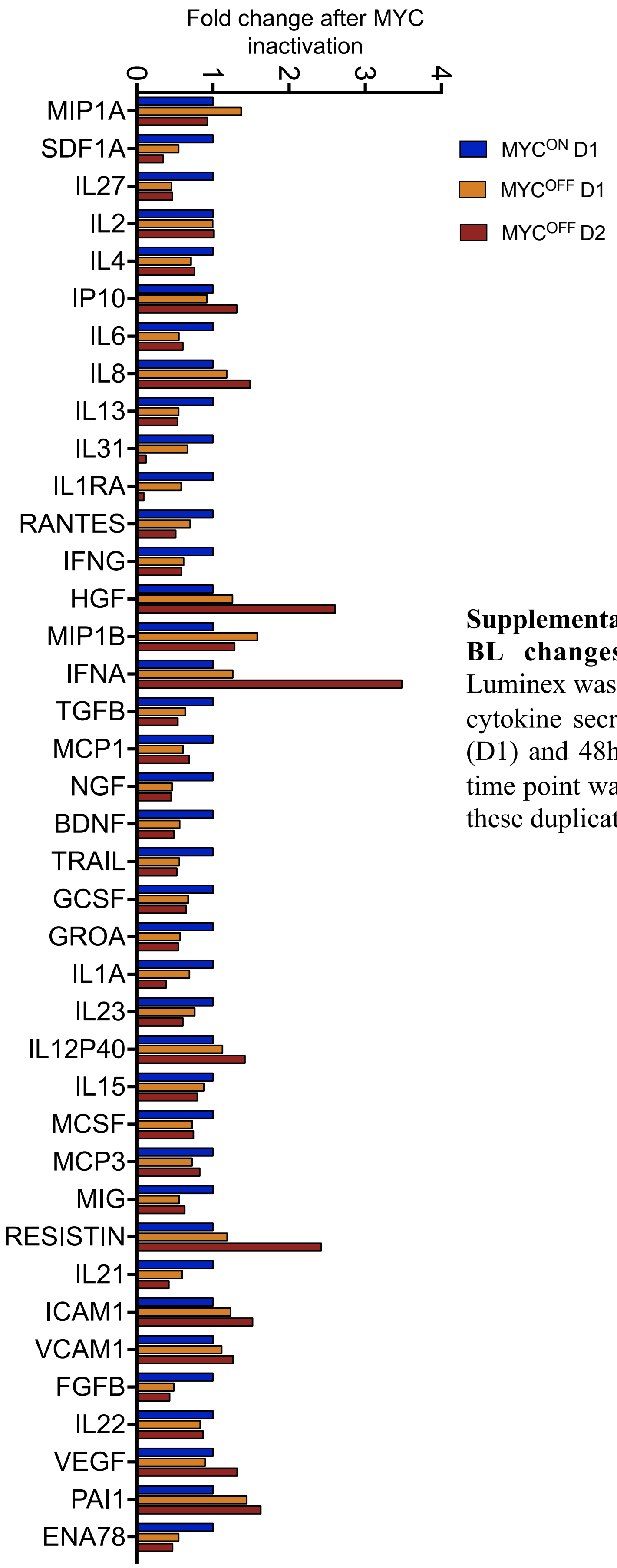


**Supplementary Figure 13: MYC inactivation in BL changes the secreted cytokine profile.** Luminex was used to evaluate the global changes in cytokine secretion post MYC inactivation for 24h (D1) and 48h (D2) in the P493-6 BL model. Each time point was carried out in duplicates. Average of these duplicates is shown.

**Supplementary Figure 14:** *Flow cytometry validation of T-lymphoma cell line derived from an SRα-tTA/Tet-O-MYC mouse*

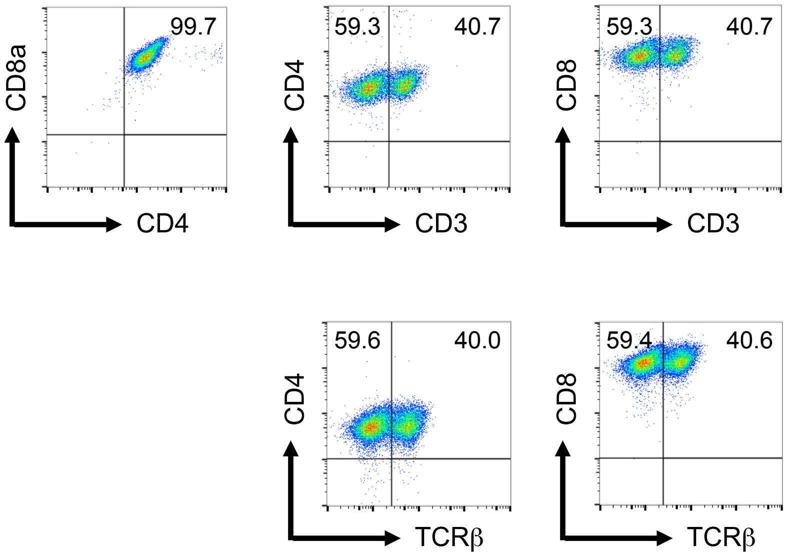


**Supplementary Figure 14: Flow cytometry validation of a T-lymphoma cell line derived from an *SRα-tTA/Tet-O-MYC* mouse.** Flow cytometry plots depicting the surface expression of T cell markers, such as CD3, CD4, CD8 and TCRβ in a cell line derived from *SRα-tTA Tet O MYC* mice.-

**Supplementary Figure 15:** *Suppression of STAT1/2-Type I IFN signaling in MYC-driven T cell lymphomas*

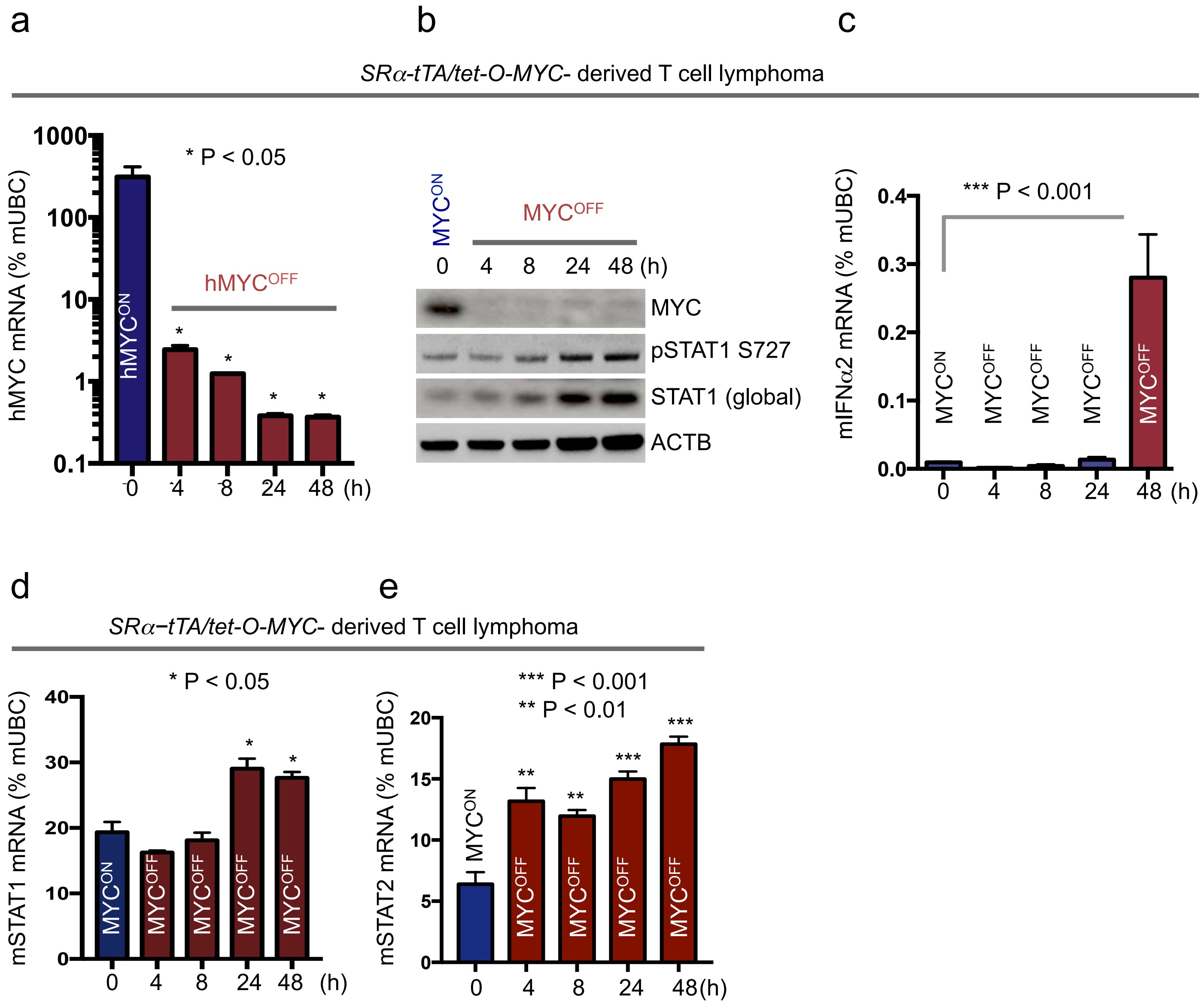


**Supplementary Figure 15: Suppression of STAT1/2-Type I IFN signaling in MYC-driven T cell lymphomas.** (**a**) Quantitative real time PCR for human MYC transgene in SRα-tTA MYC transgenic mouse T-lymphoma cells before and after MYC inhibition by doxycycline for 4, 8, 24 and 48h (n = 3, mean ± s.d). (**b**) Immunoblotting to measure changes in STAT1 activation pre- and post-MYC transgene inactivation in SRα-tTA MYC mouse T-lymphoma cells. (**c-e**) Quantitative real time PCR for mType I IFNα2 (c), mSTAT1 (d) and, mSTAT2 (e) in a T-lymphoma cell line derived from an *SRα-tTA/tet-O-MYC* mouse before and after MYC inactivation by doxycycline for 4, 8, 24 and 48 hours (n = 3, mean ± s.d). P-values have been calculated using the Student's t-test (*p < 0.05, **p < 0.01, ***p < 0.001).

**Supplementary Figure 16:** *MYC binds to STAT1 promoter in MYC-driven lymphomas*

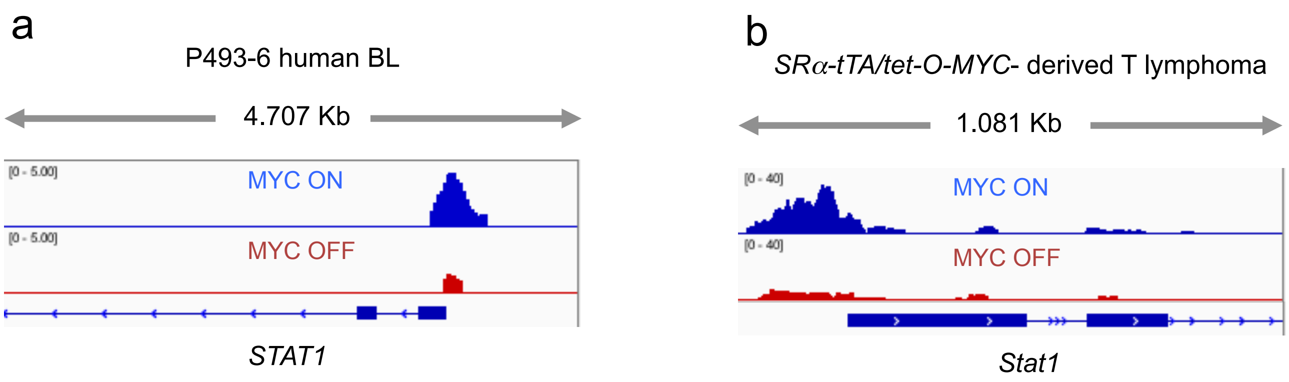


**Supplementary Figure 16: MYC binds to STAT1 promoter in MYC-driven lymphomas. (a-b)** ChIP sequencing depicting binding of MYC to STAT1 promoter in P493-6 (a. GSE36354) and SRα-tTA-MYC T lymphoma (b, GSE44672) cell lines before and after MYC inactivation.

**Supplementary Figure 17:** *Oncogenic MYC transcriptionally represses STAT1/2-Type I IFN signaling*

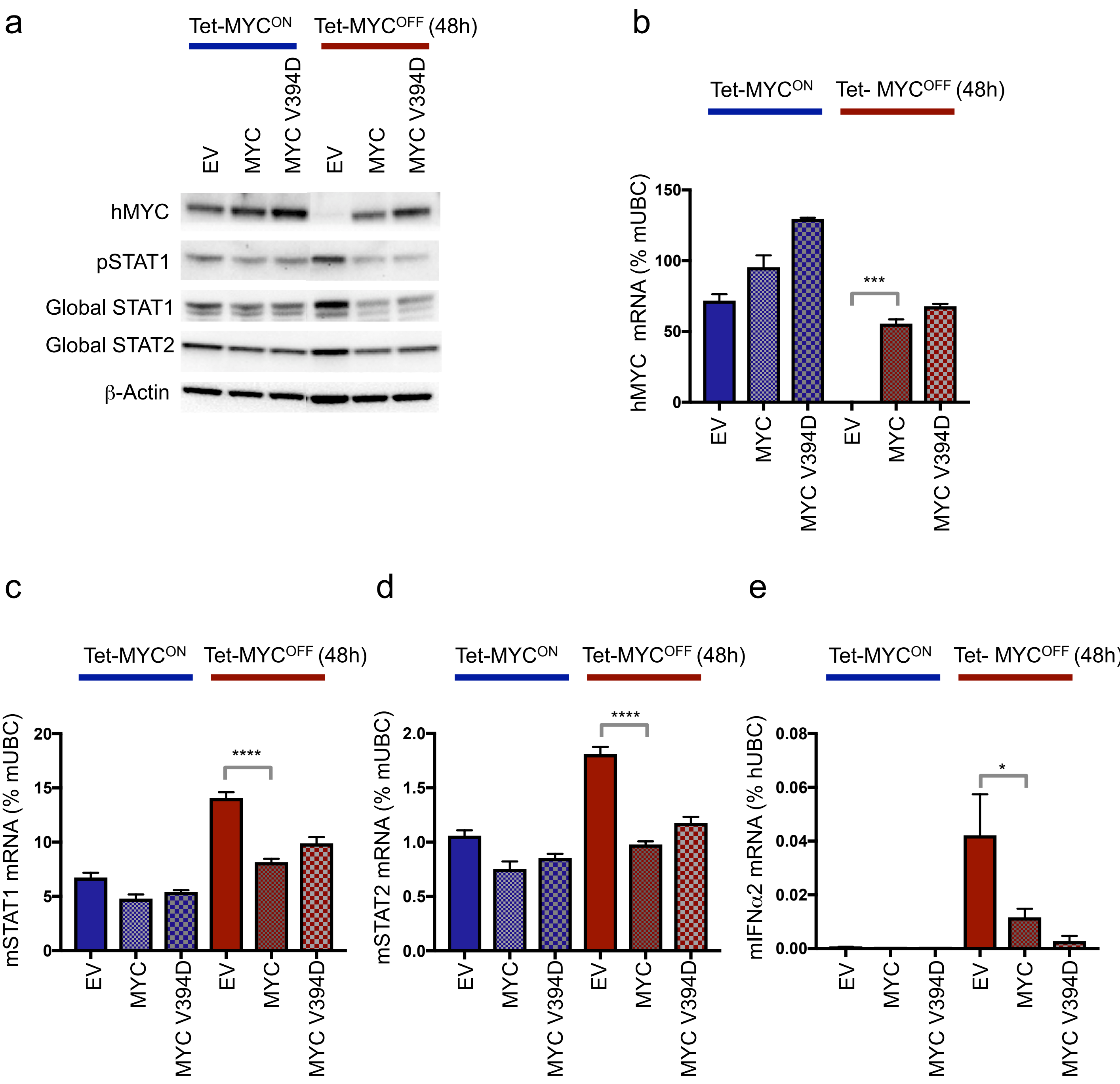


**Supplementary Figure 17: Oncogenic MYC transcriptionally represses STAT1/2-Type I IFN signaling.** (**a**) Immunoblotting in MYC-driven SRα-tTA T-lymphoma cell lines expressing empty vector (EV), MYC overexpression vector (MYC), or a MYC mutant that fails to bind MIZ1 (MYC V394D) before (Tg-MYC$^{ON}$) and after (Tg-MYC$^{OFF}$) MYC inactivation. **(b-e)** Quantitative real time PCR for hMYC (b), mSTAT1 (c), mSTAT2 (d), and mIFNα2 (e) in MYC-driven SRα-tTA T-lymphoma cell lines expressing empty vector (EV), MYC overexpression vector (MYC), or a MYC mutant that fails to bind MIZ1 (MYC V394D) before (Tg-MYC$^{ON}$) and after (Tg-MYC$^{OFF}$) MYC inactivation (n = 3, mean ± s.d). All p values have been calculated using student t-test. *$p < 0.05$; ***$p < 0.001$; ****$p < 0.0001$.

**Supplementary Figure 18:** *Isolation and purification of syngeneic NK cells for adoptive transfer into MYC-driven T-lymphoma bearing NSG recipients*

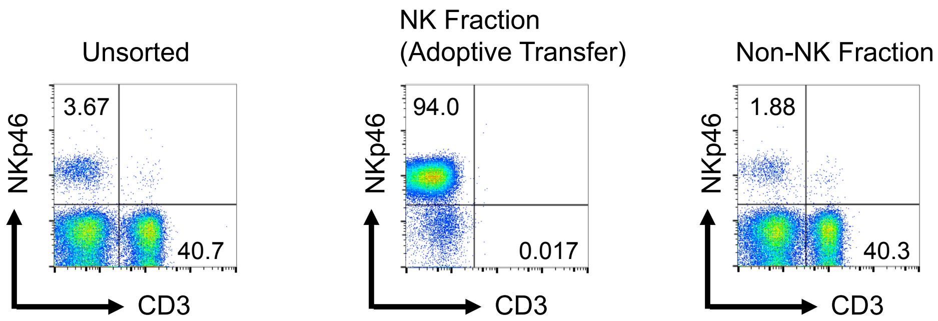


**Supplementary Figure 18: Isolation and purification of syngeneic NK cells for adoptive transfer into MYC-driven T-lymphoma bearing NSG recipients.** Validation of purity of NK cells isolated by magnetic activated cell sorting (MACS) by flow cytometry for surface CD3, NKp46 and CD49b.

**Supplementary Figure 19:** *Oncogenic MYC suppresses NK cell-mediated immune surveillance of lymphoid malignancies.*

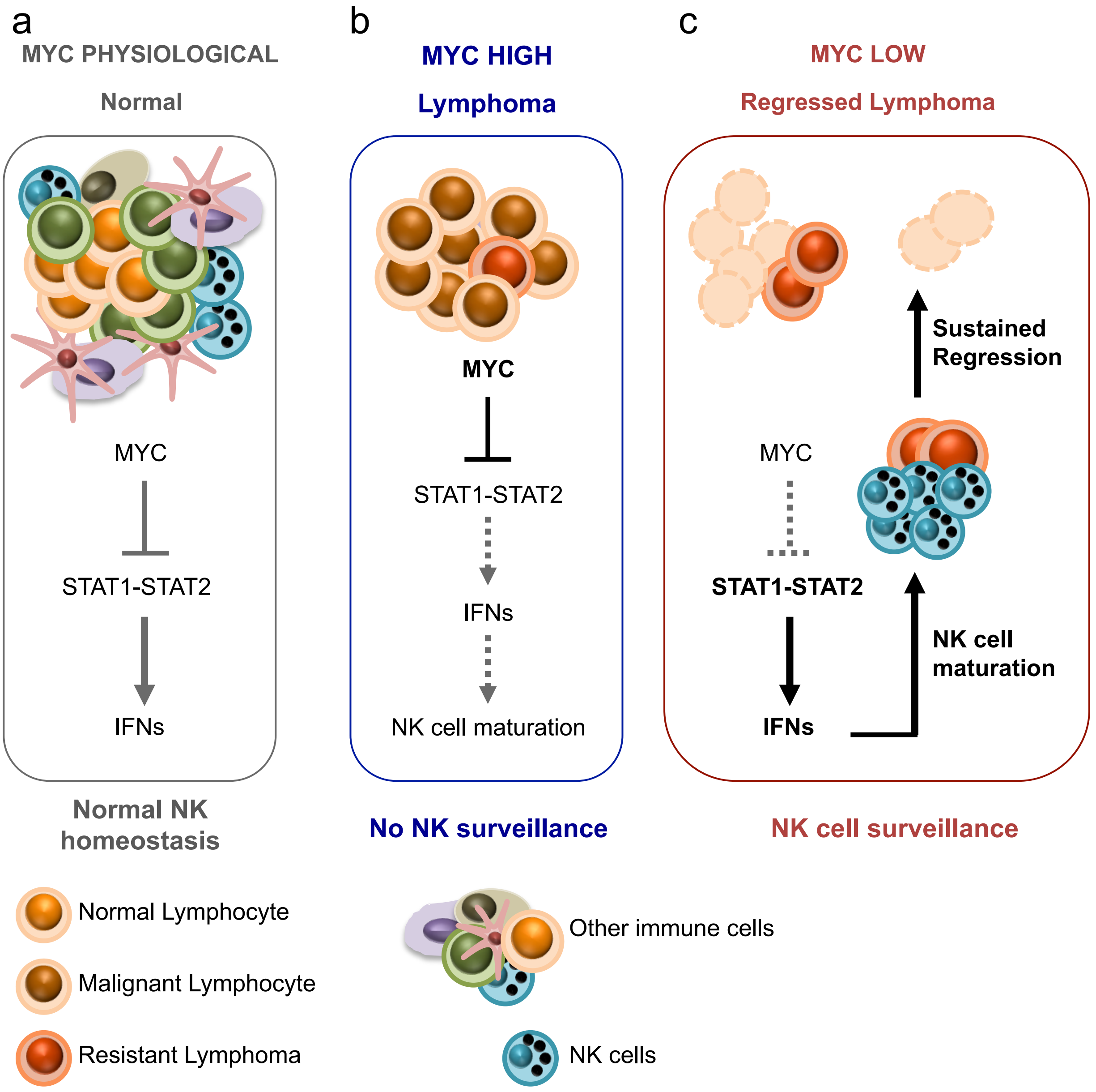


**Supplementary Figure 19: Oncogenic MYC suppresses NK cell-mediated immune surveillance of lymphoid malignancies.** (**a**) Normal distribution of splenic immune compartments in the absence of lymphoma. (**b**) MYC-induced lymphomagenesis is associated with the modulation of cytokines and immune subsets in the tumor microenvironment. Activation of oncogenic MYC reduces the secretion of anti-lymphomagenic cytokines, such as Type I IFNs. The altered cytokine milieu excludes NK cells reducing immune surveillance of the lymphoma. (**c**) MYC inactivation triggers Type I IFN secretion by the lymphoma and NK-mediated immune surveillance. Hyperactivated signaling arms under each condition are depicted in bold black.

**Supplementary Figure 20:** *Gating strategy for CyTOF and flow cytometry*

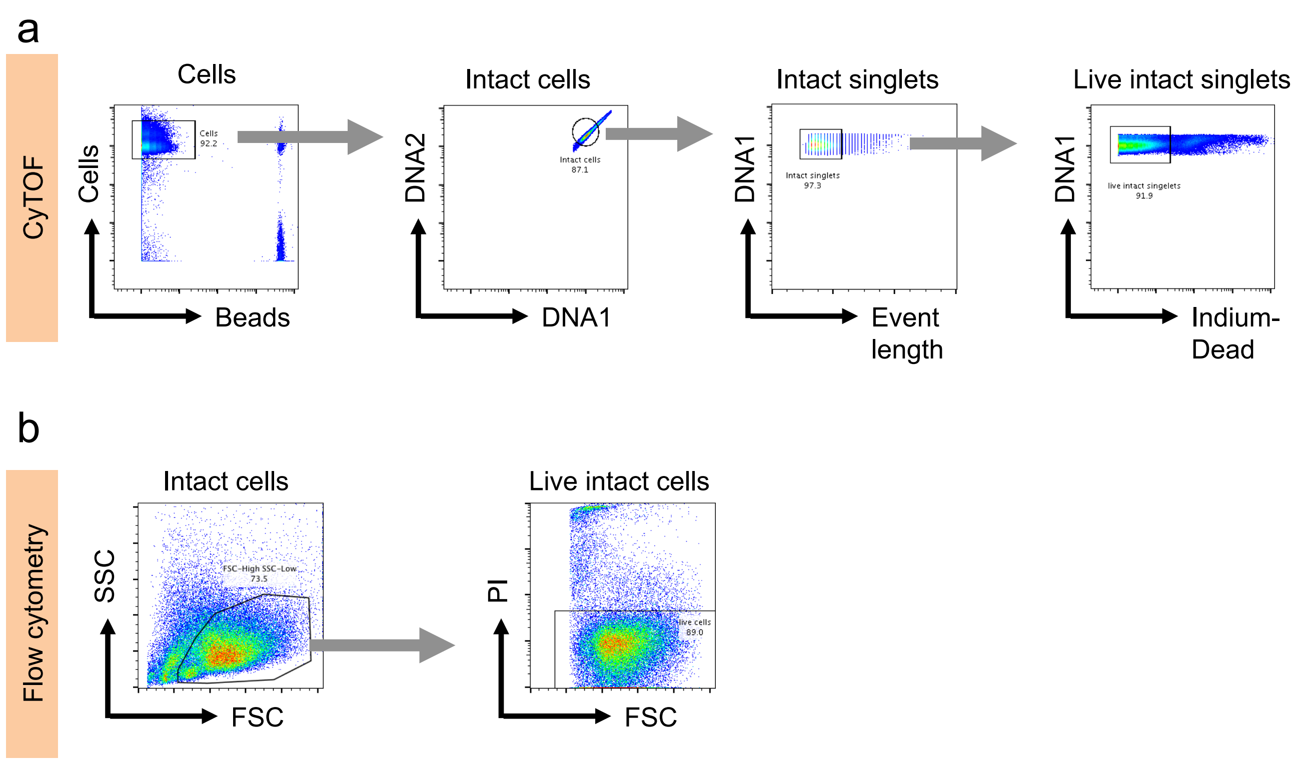

**Supplementary Table 1:** *Antibodies used for CyTOF immunophenotyping (mouse)*

| Antigen | Metal Tag | Source |
|---|---|---|
| Live-Dead | In115 | HIMC, Stanford University |
| Ly6G | Pr141 | Fluidigm Corporation |
| CD11c | Nd142 | Fluidigm Corporation |
| CD11b | Nd143 | Fluidigm Corporation |
| Ly6C | Dy162 | Fluidigm Corporation |
| CD3 | Sm152 | Fluidigm Corporation |
| CD8a | Er168 | Fluidigm Corporation |
| CD4 | Nd145 | Fluidigm Corporation |
| TCRγδ | Tb159 | Fluidigm Corporation |
| TCRβ | Tm169 | Fluidigm Corporation |
| CD19 | Sm149 | Fluidigm Corporation |
| MHCII | Yb174 | Fluidigm Corporation |
| NKp46 | Eu153 | Fluidigm Corporation |
| CD49b | Er170 | Fluidigm Corporation |
| Ter119 | Sm154 | Fluidigm Corporation |

**Supplementary Table 2:** *Antibodies used for Phospho-CyTOF (human)*

| Antigen | Metal Tag |
|---|---|
| Live-Dead | In115 |
| Ly6G | Pr141 |
| CD19 | Nd142 |
| CD117 | Nd143 |
| CD11b | Nd144 |
| CD4 | Nd145 |
| CD20 | Sm147 |
| CD7/ CD45RO | Sm149 |
| CD123 | Eu151 |
| CD27 | Sm152 |
| CD45RA | Eu153 |
| CD45 | Sm154 |
| pP38 | Gd156 |
| CD24 | Gd157 |
| pSTAT3 | Gd158 |
| CD11c | Tb159 |
| CD14 | Gd160 |
| IgD | Dy161 |
| pERK1/2 | Dy162 |
| IκBtot | Dy163 |
| CD25 | Dy164 |
| pS6 | Ho165 |
| CD16 | Er166 |
| CD38 | Er167 |
| CD8 | Er168 |
| pSTAT1 | Tm169 |
| CD3 | Er170 |
| pSTAT5 | Yb172 |
| pPLCγ2 | Yb173 |
| HLADR | Yb174 |
| CD56 | Lu175 |
| CD127 | Yb176 |

**Supplementary Table 3:** *Antibodies used for flow cytometry*

| Surface Antigen | Clone ID/ Catalog No. | Reactivity | Source |
|---|---|---|---|
| CD3e | 145-2C11 | Mouse | BD Pharmingen |
| CD4 | RM4-5 | Mouse | BD Pharmingen |
| CD8a | 53-6.7 | Mouse | BD Pharmingen |
| CD335 (NKp46) | 29A1.4 | Mouse | BioLegend |
| CD122 | TM-beta1 | Mouse | BD Pharmingen |
| CD16/CD32 (Fc block) | 2.4G2 | Mouse | BD Pharmingen |

**Supplementary Table 4:** *Antibodies used for immunoblotting*

| Antigen | Clone ID | Dilution | Source |
|---|---|---|---|
| MYC | N-262 | 1:1000 | Santa Cruz Biotechnology |
| ACTB | ab8227 | 1:10,000 | Abcam |
| STAT1 | 9172 | 1:1000 | Cell Signaling Technology |
| Phospho S727 STAT1 | 9177 | 1:1000 | Cell Signaling Technology |
| STAT2 | D9J7L | 1:1000 | Cell Signaling Technology |

**Supplementary Table 5:** *Lentiviral and retroviral vectors used in the study*

| Vector | Type | Source |
|---|---|---|
| pMSCV Luciferase IRES-puro | Retroviral | Addgene |
| pRRL | Lentiviral | Lab of Martin Eilers |
| pRRL hMYC | Lentiviral | Lab of Martin Eilers |
| pRRL hMYC V394D | Lentiviral | Lab of Martin Eilers |

Transfections of the above retroviral constructs were performed as discussed in the online methods section.

**Supplementary Table 6:** *Sequences of oligonucleotide primers used for quantitative PCR*

**Mouse primers**

| | |
|---|---|
| mIFNA2_F | 5’-AAGGTCCTGGCACAGATGAG-3’ |
| mIFNA2_R | 5’-GGAGGGTTGTATTCCAAGCA-3’ |
| mSTAT1_F | 5’-TGGTGAAATTGCAAGAGCTG-3’ |
| mSTAT1_R | 5’-CAGACTTCCGTTGGTGGATT-3’ |
| mSTAT2_F | 5’-GCCTCCATTCTCTGGTTCAA-3’ |
| mSTAT2_R | 5’-TCCTCCATCTTGCAGCTCTT-3’ |
| mUBC_F | 5’-AGCCCAGTGTTACCACCAAG-3’ |
| mUBC_R | 5’-ACCCAAGAACAAGCACAAGG-3’ |

**Human primers**

| | |
|---|---|
| hIFNA2_F | 5’-GCAAGTCAAGCTGCTCTGTG-3’ |
| hIFNA2_R | 5’-CAAACTCCTCCTGGGGAAAT-3’ |
| hSTAT1_F | 5’-CCGTTTTCATGACCTCCTGT-3’ |
| hSTAT1_R | 5’-TGAATATTCCCCGACTGAGC-3’ |
| hSTAT2_F | 5’-GGAACAGCTGGAGACATGGT-3’ |
| hSTAT2_R | 5’-GGCTGGGTTTCTACCACAAA-3’ |
| hMYC_F | 5’-CTGCGACGAGGAGGAGAACT-3’ |
| hMYC_R | 5’-GGCAGCAGCTCGAATTTCTT-3’ |
| hUBC_F | 5’-CTGGAAGATGGTCGTACCCTG-3’ |
| hUBC_R | 5’-GGTCTTGCCAGTGAGTGTCT-3’ |
